# Species existence and coexistence under nutrient enrichment in the Park Grass

**DOI:** 10.1101/2022.06.08.495203

**Authors:** Qi Yao, Yanhao Feng

**Author notes:** Yanhao Feng, **Email:**. **Author Contributions:** YHF conceived and designed the research, QY performed the research and wrote the first draft of the manuscript, and both authors contributed substantially to revisions. **Data sharing plans:** All the data used in this study are publicly available from the electronic Rothamsted Archive (e-RA) (www.era.rothamsted.ac.uk), which is a permanent online source for data from the Rothamsted long-term experiments (https://www.nature.com/articles/sdata201872). We will provide R code to download and organize the data from the e-RA for reproducing our results. All the R code associated with this study will be archived in Zenodo.

## Abstract

Numerous studies have shown that nutrient enrichment causes the loss of plant diversity in different grasslands across the globe. Thus far, three main hypotheses (niche dimension, competitive asymmetry, and soil acidification) have been proposed to account for this general phenomenon, but our knowledge of the underlying mechanisms remains rather vague. To reveal the cryptic mechanisms, we analyzed the famous long-term Park Grass Experiment (1856-) under modern coexistence theory by fitting Lotka-Volterra competition models with time-series data from the different treatments (15 different combinations of nutrient addition fully crossed with four levels of soil pH) to quantitatively test the three competing hypotheses. Supportive of the competitive asymmetry and soil acidification hypotheses, both nutrient addition and soil acidification overall decreased intrinsic population growth rates (*r*) and intensified competitive differences or asymmetries dramatically, which mostly favored grasses over forbs and legumes (and also forbs over legumes). These changes in *r* and competitive differences or asymmetries are generally consistent with the abundance changes of different functional groups following the various treatments. Moreover, the altered *r* (determining species *existence*) and competitive differences (affecting species *coexistence*) effectively explained the diversity loss and recovery (after nitrogen addition was withheld). However, while nutrient addition significantly decreased per-capita intra- and inter-specific competition (which indicates that belowground competition becomes less intense when soil nutrients are more abundant), it did not decrease niche differences as predicted, poorly supporting the niche dimension hypothesis. These findings advance our understanding of fundamental mechanisms driving the response of plant communities to nutrient deposition in nature.

**Significance Statement:** An unresolved fundamental scientific mystery with the crucial applied value in ecology is what causes the general loss of biodiversity following nutrient enrichment in the Anthropocene. In this study, we combined the recent advance in coexistence theory with the longest-running fertilization experiment existing in the world to wrestle with this conundrum. Our major results, which highlight the critical role of both existence and coexistence, could help settle the debate between the three popular hypotheses and also, for the first time, provide quantitative explanations for the general findings in numerous nutrient-addition experiments. Our study shows the importance of applying modern coexistence theory to more quantitatively explain, predict and cope with the responses of ecological communities to global change factors in nature.

## Introduction

Along with accelerating anthropogenic activities across the globe, nutrient deposition has increasingly posed one of the major threats to the biodiversity, structure and stability of natural ecosystems (1-3). For example, numerous studies have shown that addition of various nutrients (e.g., N, P, K and trace elements) causes the decline of plant diversity globally in different types of grasslands (4-6). Thus far, three main hypotheses have been proposed to account for this general phenomenon, i.e., niche dimension hypothesis (7), competitive asymmetry hypothesis (8, 9) and soil acidification hypothesis (10). The niche dimension hypothesis is derived from classical coexistence theory, which asserts the importance of multiple limiting factors (e.g., resources, temperature, space) for maintaining species diversity (11-14). Specifically, the hypothesis proposes that the decline of plant diversity is because the dimensions of the “hypervolume” niche space consisting of multiple limiting resources (in which species differentiate to enable coexistence, *sensu* Hutchinson 1957) could reduce along with nutrient addition (7). More precisely, it is because nutrient addition is supposed to change the corresponding limiting resources into non-limiting ones, leading to the removal of niche axes (i.e., dimensions). Although this hypothesis is theoretically self-consistent, the challenge is to test it empirically because of the difficulty in quantifying niche per se and its dimensions in nature (15, 16). Therefore, the empirical tests hitherto are largely indirect without rigorously quantifying the reduction of niche dimensions (but using the number of added nutrients as a surrogate instead) (4, 7), which makes the hypothesis yet in debate (17, 18).

The competitive asymmetry hypothesis, in contrast, claims that the intensified asymmetry in competition (for light or nutrients) drives the decline of plant diversity under nutrient enrichment (8, 9, 19). Specifically, nutrient addition is supposed to decrease the belowground competition for soil nutrients (often considered symmetric, but see 20) but increase the intensity and asymmetry of aboveground competition for light with unidirectional nature (21, 22). As a result, the increasingly taller and larger species under nutrient addition could pre-empt a more disproportionate fraction of light (i.e., size-asymmetric competition) and thereby drive the smaller species locally extinct (8, 19, 23). Please note there is also evidence that nutrient addition could affect the degree of asymmetry in belowground competition for soil nutrients (9, 24). As opposed to the hypotheses from a resource perspective, the soil acidification hypothesis stresses the role of soil pH decrease often concomitant with N addition in accelerating species loss (10, 25). As an edaphic stress, soil acidification reduces plant diversity because it could inhibit plant growth and survival by increasing metal toxicity, causing cation leaching and/or lowering nutrient availability (26). Taken together, despite the experimental tests for the three competing hypotheses, poorly known is how the purported mechanisms strictly translate into population growth and dynamics and species interactions to cause community diversity loss. In other words, no research has quantitatively evaluated the three hypotheses, which makes our knowledge of the underlying mechanisms remain rather vague.

Modern coexistence theory proposes a feasible framework to define and quantify niche and competitive differences between species (27, 28). In essence, niche difference measures the degree to which a species limits itself more than its competitor, while competitive difference (i.e., fitness difference, *sensu* Chesson 2000) quantifies the overall asymmetry in competitive ability between species (29). In theory, niche difference favors coexistence through “stabilising” mechanisms, as it generates negative density-dependent regulations which enable a species to recover when rare. In contrast, competitive difference drives exclusion through density-independent processes, and coexistence can be fostered by equalizing this difference (“equalizing” mechanisms). Mathematically, niche and competitive differences are often calculated with parameters in dynamic competition models (e.g., Lotka-Volterra competition model, see *Theory* for details), which can be fitted with empirical data to understand species coexistence in nature (30-33). Importantly, niche and competitive differences as emergent properties are rooted in biological mechanisms of species coexistence (27, 34). Specifically, niche differences emerge from species’ differentiation (or trade-offs) in multiple limiting factors (e.g., resources) (14, 35). In contrast, competitive differences arise from any biological process determining the overall difference between species in their abilities to compete for resources (e.g., nutrients, light, water) (29).

Thus, modern coexistence theory could provide a powerful toolbox to quantitatively test the three hypotheses. Specifically, niche difference should decrease accordingly if niche dimensions are reduced, and competitive difference should effectively capture the intensified competitive asymmetry. Following this, we infer that nutrient addition causes plant diversity loss possibly because it could promote competitive exclusion by decreasing niche differences (niche dimension hypothesis), intensifying competitive differences (competitive asymmetry hypothesis) or both. Besides, soil acidification could interact with nutrient addition to accelerate species loss by affecting niche and competitive differences (soil acidification hypothesis). Lastly, in addition to the failure of species *coexistence* as we have often focused on, the loss of plant diversity could also occur if species *existence* is jeopardized. For instance, the mortality of some species could increase with nutrient addition increasing light asymmetry (19) and soil acidification (10) (competitive asymmetry and soil acidification hypotheses). This should quantitatively translate into the plunge of intrinsic population growth rates (*r*) (36), driving the species to vanish.

To quantitively re-evaluate the three hypotheses and reveal the cryptic mechanisms, we analyzed the famous long-term Park Grass Experiment (1856-) under modern coexistence theory by fitting Lotka-Volterra competition models with time-series data. Briefly, we found clear quantitative evidence supportive of the competitive asymmetry and soil acidification hypotheses but not the niche dimension hypothesis. Specifically, our results showed that nutrient addition and soil acidification reduced the plant diversity by decreasing *r* (determining species *existence*) and intensifying competitive differences (affecting species *coexistence*) without decreasing niche differences. These novel findings explicitly elucidate the relative importance of the three hypotheses and also greatly deepen our insights into the fundamental mechanisms driving the decline of plant diversity under nutrient enrichment.

## Results

### Effects of nutrient addition and soil pH on r and niche and competitive differences

For the detailed statistical results, see Table S4. Overall, *r* decreased with addition of N (more for NH_4_^+^ than NO_3_^-^; Fig. 1a) and NaMg (Fig. S5.1b) but increased with addition of P (Fig. 1b) and K (Fig. S5.1a). Accordingly, *r* rebounded after N addition was withheld, more for N*2PKNaMg (NO_3_^-^) than N2PKNaMg (NH_4_^+^) (Fig. 4a). In addition, the effects of different nutrients interacted. Specifically, P addition increased *r* when NO_3_^-^ or K is present or N is absent (Figs 1d-e) but decreased *r* when NH_4_^+^ is present (Fig. 1d). Moreover, K addition increased *r* when P is present but decreased *r* when P is absent (Fig. 1e). More importantly, the effects of nutrient addition differed for three functional groups. For example, N addition overall decreased *r* more for legumes and grasses than forbs (Fig. 1g) (N addition × functional group interaction: χ^2^=6.72, P=0.034), while P addition increased *r* more for legumes than grasses and forbs (Fig. 1h). Accordingly, *r* rebounded more for legumes and grasses than forbs after withholding N addition (Fig. 4b). Lastly, *r* decreased dramatically as soil pH decreased (Fig. 1c) and the effects were stronger when NH_4_^+^ (rather than NO_3_^-^) (Fig. 1f) or P (Fig. S5.1d) is present, indicating that nutrient addition and soil acidification are synergistic. Moreover, soil acidification decreased *r* more for legumes than grasses and forbs (Fig. 1f).

**Figure 1.**
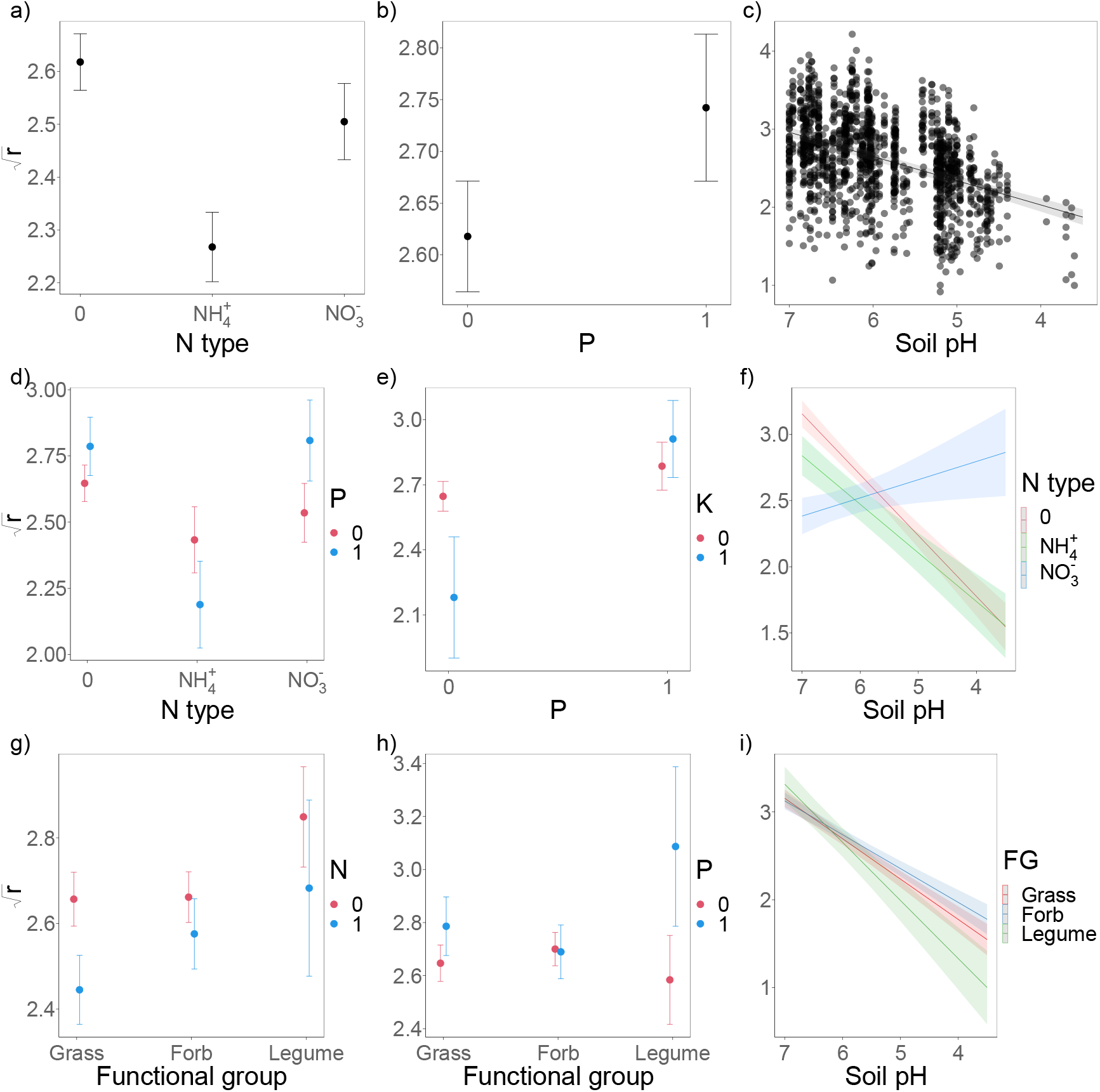
The effects of a) N type (0, NH_4_^+^, NO_3_^-^), b) P (0, 1) and c) soil pH, the interactions between d) N type and P, e) P and K (0, 1), and f) N type and soil pH, and the interactions between functional group (FG: grass, forb and legume) and g) N (0, 1), h) P, and i) soil pH, on intrinsic population growth rate (*r*). The lines, means (circles) and standard errors (error bars) are predicted from the statistical models (see Table S4 in SI Appendix S3 for the detailed results).

Both niche and competitive differences changed considerably in response to the various treatments of nutrient addition and soil pH (Fig. 2). However, niche differences overall did not decrease or even increased with nutrient addition (Fig. 3a; Table S4 and Fig. S5.2) and also increased as soil pH decreased (Fig. 3b; Fig. S5.2d). In contrast, nutrient addition and soil acidification intensified competitive differences or asymmetries significantly and synergistically (Table S4 and Table S6). Overall, addition of N or P and soil acidification both made grasses more competitive than forbs and legumes, and also forbs more competitive than legumes (Figs 3d and 3g; Fig. S5.3g). However, adding K (or NaMg) made legumes more competitive than grasses and forbs (Figs S5.3h-i).

**Figure 2.**
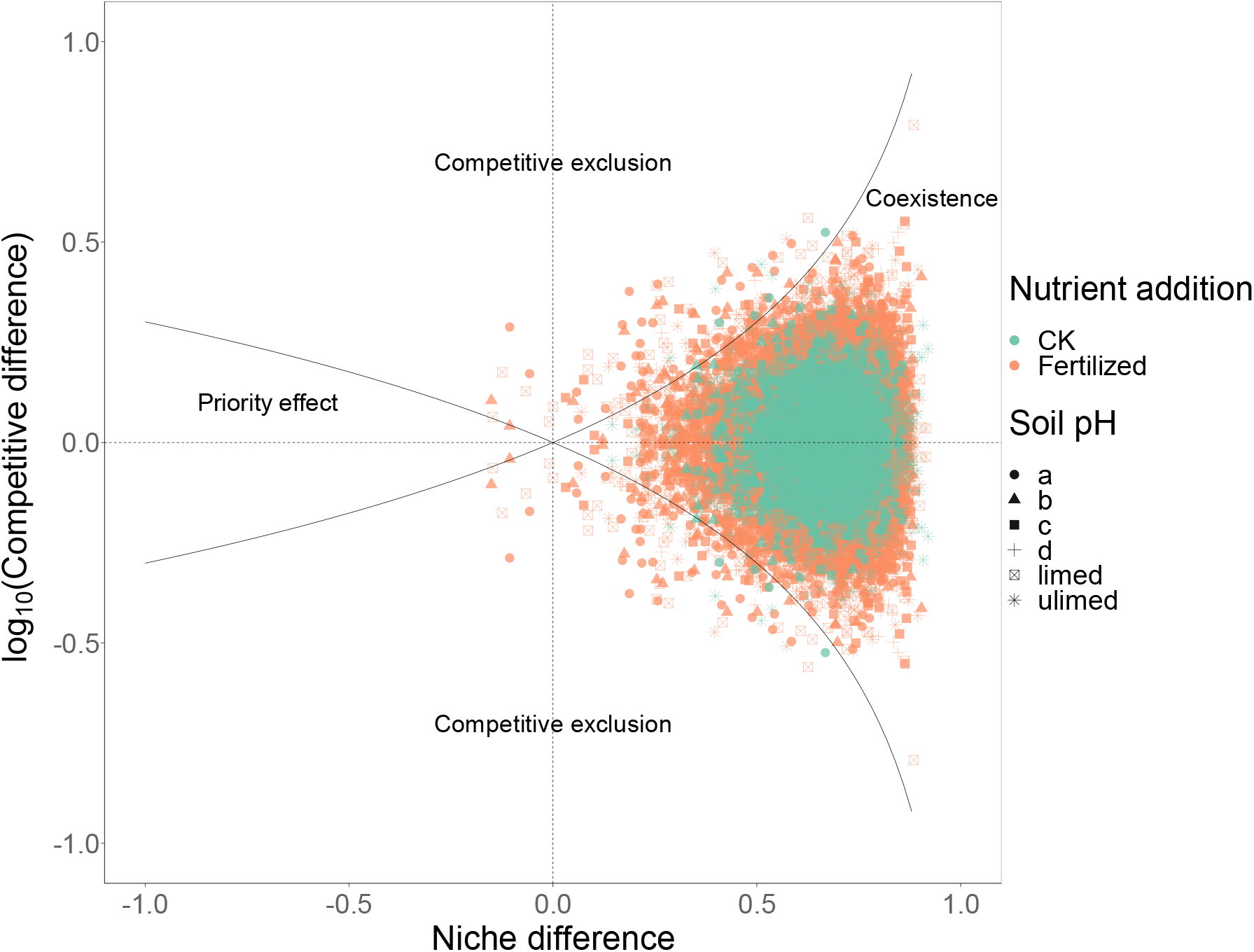
The responses of niche and competitive differences to all treatments of nutrient addition (CK: without fertilization; fertilized: all fertilized treatments) and soil pH. There are in total 11455 pairs of niche and competitive differences (CK: 2883 pairs; fertilized: 8572 pairs). The “a”, “b”, “c” and “d” indicate quarter-plots with four different levels of soil pH treatments (soil pH: 7, 6, 5 and unlimed, respectively) in the “recent” subset (1965-2000). The “limed” and “unlimed” indicate the limed and unlimed half-plots respectively in the “longul” subset (1856-2000). The areas of stable coexistence, competitive exclusion and priority effect are also indicated. More details about the treatments can be found in SI Appendix S1.

Interestingly, both per-capita intraspecific and interspecific competition (*a*_*ii*_ and *a*_*ij*_) decreased dramatically with nutrient addition (Fig. 3c; Figs S5.4a-c and S5.5a-c for individual nutrients) but increased with soil acidification (Figs S5.4d and S5.5d). Conversely, both *a*_*ii*_ and *a*_*ij*_ increased after withholding N addition (Fig. 4c). More importantly, the effects of both nutrient addition and soil acidification on *a*_*ii*_ or *a*_*ij*_ varied for different functional groups. Specifically, *a*_*ii*_ of grasses or forbs decreased (i.e., carrying capacity, *K*_*i*_, increased) more than those of legumes (whose *a*_*ii*_ in fact increased, i.e., *K*_*i*_ decreased) with N addition (Fig. 3e). Accordingly, *a*_*ij*_ of grasses or forbs on legumes also decreased more than the reverse (*a*_*ij*_ of legumes on grasses or forbs in fact increased) with N addition (Fig. 3f) and withholding N addition tended to reverse the effects (Fig. 4d). However, K addition decreased *a*_*ii*_ (i.e., increased *K*_*i*_) more for legumes than grasses and forbs (Fig. S5.4h), and accordingly K (or NaMg) addition decreased *a*_*ij*_ of legumes on grasses and forbs more than the reverse (Figs S5.4k-l). Moreover, soil acidification increased *a*_*ii*_ (i.e., decreased *K*_*i*_) less for grasses than forbs and legumes (and less for forbs than legumes) (Fig. 3h), and accordingly increased *a*_*ij*_ of grasses on forbs and legumes (and *a*_*ij*_ of forbs on legumes) less than the reverse (Fig. 3i). Thus, a functional group became overall more competitive than another with nutrient addition or soil acidification (as mentioned above) by acquiring an advantage in *K*_*i*_ (i.e., 1/*a*_*ii*_) overriding the concomitant disadvantage in *a*_*ij*_.

**Figure 3.**
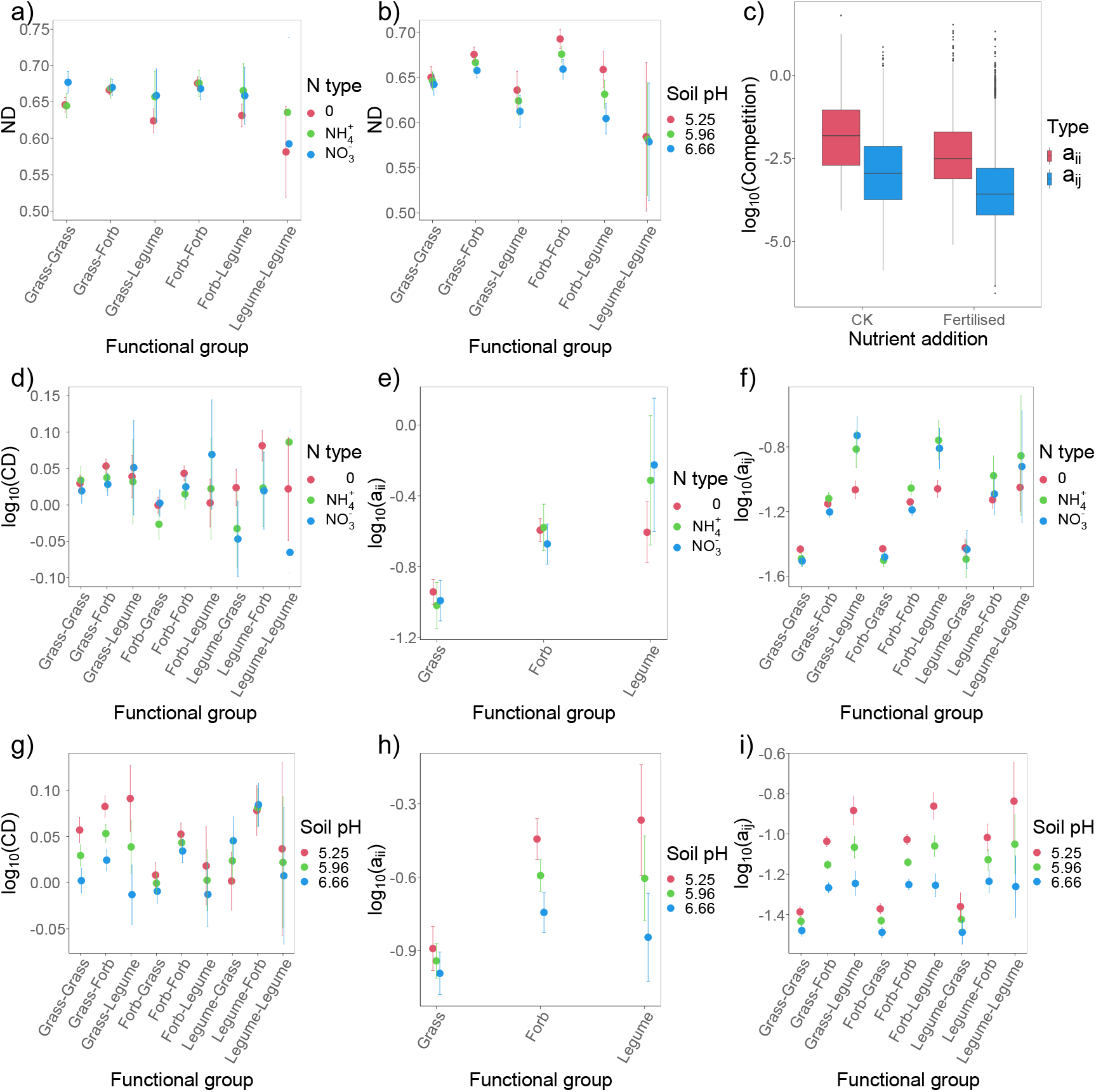
The effects of a) N type (0, NH_4_^+^, NO_3_^-^) and b) soil pH on niche difference (ND: 1–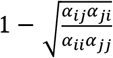 the effect of overall nutrient addition (CK: without fertilization; fertilized: all fertilized treatments) on per-capita intraspecific and interspecific competition (*a*_*ii*_ and *a*_*ij*_), and the interactions between “functional group” (for *a*_*ii*_, three functional groups: grass, forb and legume; for CD and *a*_*ij*_, all nine permutations of three functional groups) and N type or soil pH on competitive difference (CD: 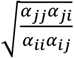; d or g), *a_ii_* (e or h) and *a_ij_* (f or i). Note that in Figs 3d and 3g, an increase in CD (functional group *i* - functional group *j*) indicates that the first functional group (*i*) becomes more competitive than the second one (*j*). Moreover, note that in Figs 3f and 3i, an increase in *a*_*ij*_ (functional group *i* - functional group *j*) indicates that the per-capita competitive effects of the second functional group (*j*) on the first one (*i*) becomes greater. The means (circles) and standard errors (error bars) are predicted from the statistical models (see Table S4 in SI Appendix S3 for the detailed results). Fig. 3c is plotted with original data but the effect is statistically significant.

**Figure 4.**
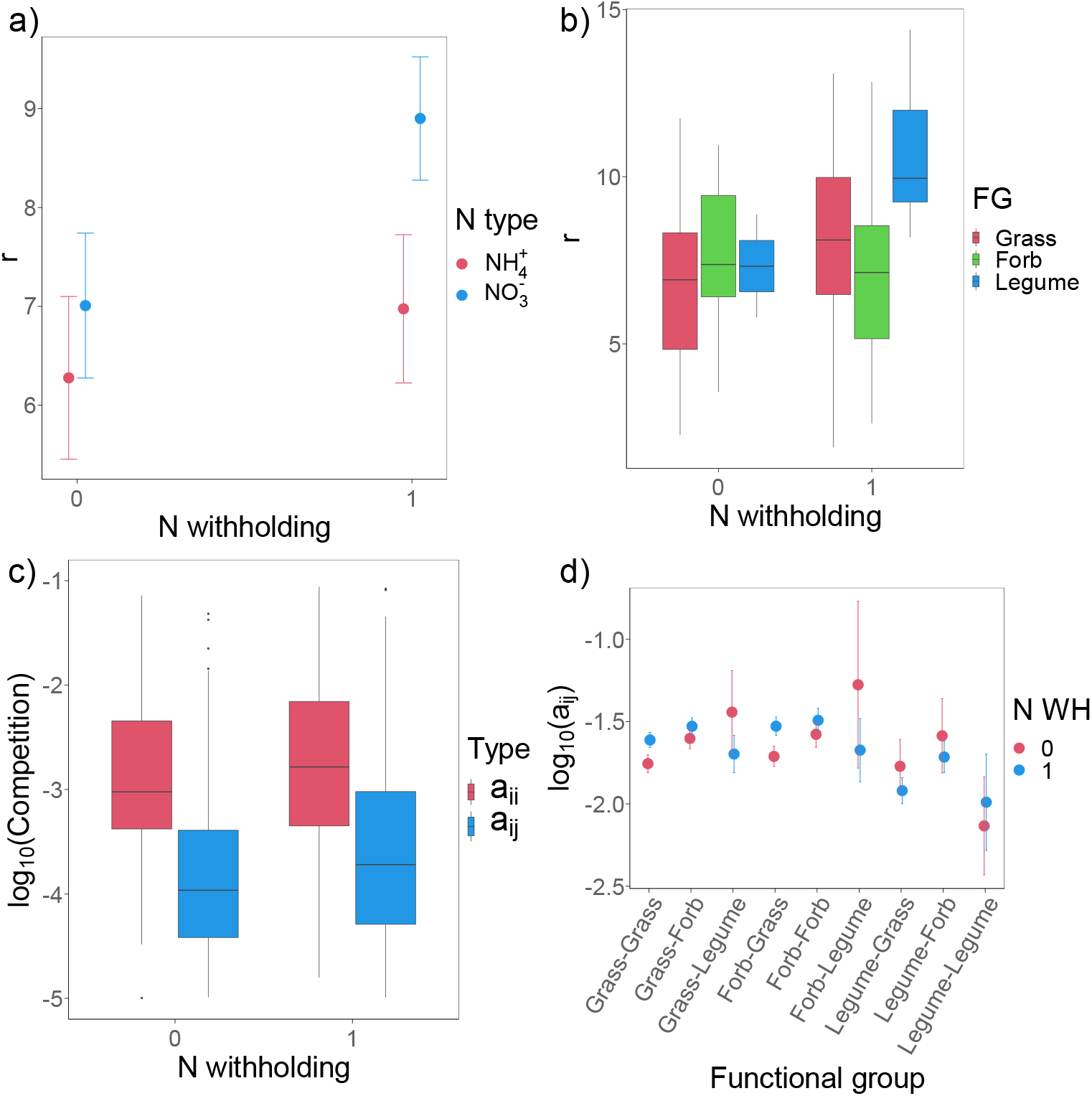
The effects of withholding N addition (N WH: 0, 1) in half of plots with the treatments of N2PKNaMg (NH_4_^+^) and N*2PKNaMg (NO_3_^-^) on intrinsic population growth rates (*r*) for a) different N types (NH_4_^+^, NO_3_^-^) and b) different functional groups (FG: grass, forb and legume), on c) per-capita intraspecific and interspecific competition (*a*_*ii*_ and *a*_*ij*_), and on d) interspecific competition (*a*_*ij*_) between different functional groups. Note that in Fig. 4d, an increase in *a*_*ij*_ (functional group *i* - functional group *j*) indicates that the per-capita competitive effects of the second functional group (*j*) on the first one (*i*) becomes greater. The means (circles) and standard errors (error bars) are predicted from the statistical models (see Table S5 in SI Appendix S3 for the detailed results). Figs 4b and 4c are plotted with original data, but the effects are all statistically significant.

### Effects of altered r and niche and competitive differences on species richness

Species richness decreased significantly as *r* decreased and as absolute competitive differences increased, but only slightly increased as niche differences increased (Figs 5a-c and Table S7 in SI Appendix S3). Moreover, *r* and niche and absolute competitive differences interacted to affect species richness (Figs 5d-f and Table S7). Specifically, the positive effect of niche differences was stronger when *r* was greater, while the negative effect of absolute competitive differences was stronger when *r* was smaller (Figs 5d-e). Moreover, the positive effect of niche differences was stronger when absolute competitive differences were greater, and conversely the negative effect of absolute competitive differences was stronger when niche difference was smaller (Fig. 5f).

**Figure 5.**
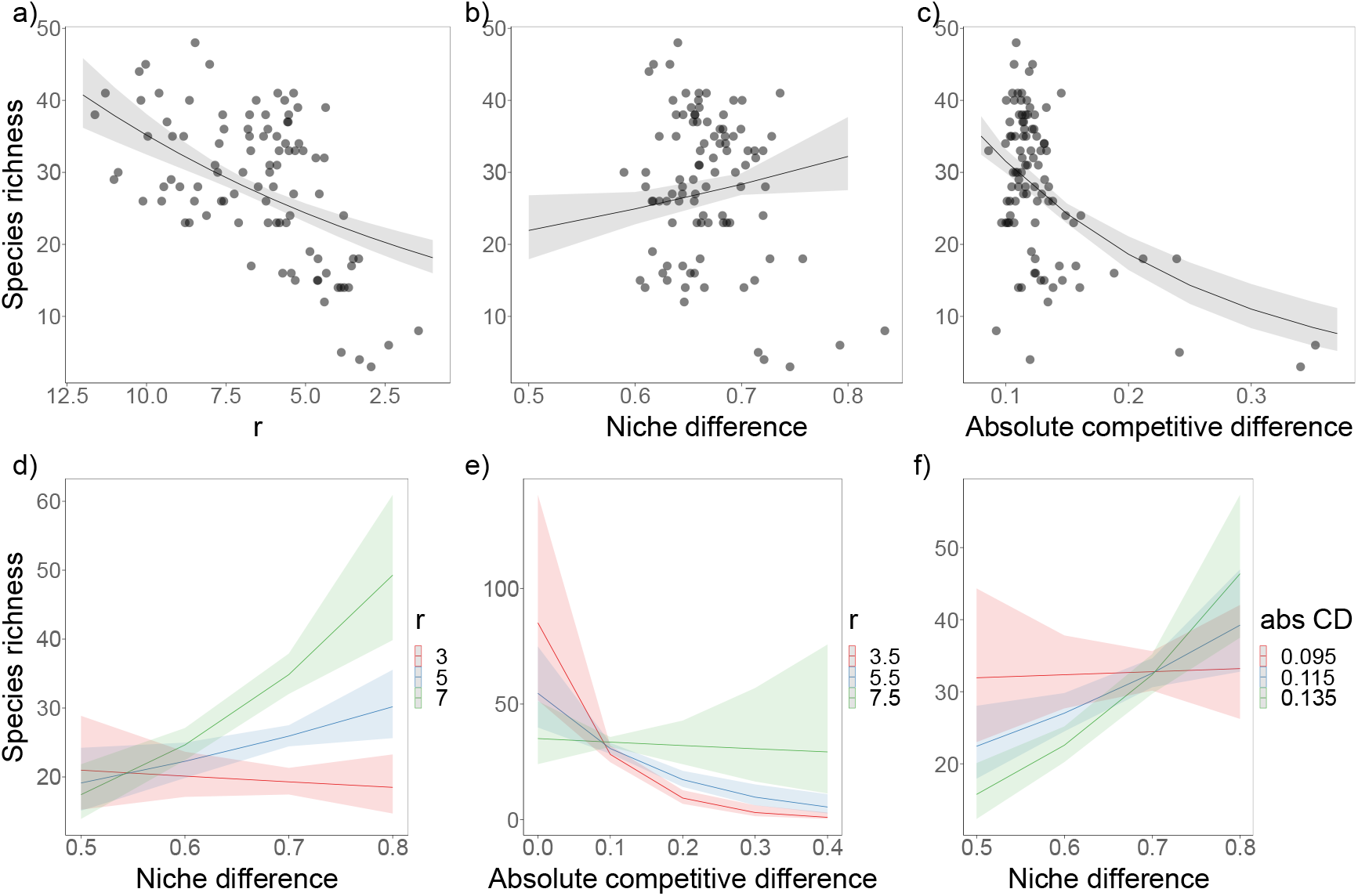
The effects of a) intrinsic population growth rate (*r*), b) niche difference, c) absolute competitive difference (abs CD), and d-f) their interactions on species richness. Note that all the response and explanatory variables are plot-level averages, and species richness is averaged from 1991 to 2000. Absolute competitive differences are absolute values of log_10_-transformed competitive differences and reflect absolute competitive asymmetries between species. The dots are original data points, and the lines are predicted from the statistical models (see Table S7 in SI Appendix S3 for the detailed results).

## Discussion

### Existence under nutrient enrichment

Existence is prior to coexistence. The competition-free *r* as a fundamental metric of population has been proposed to quantitatively examine species persistence or extinction under environmental changes (36-38). Our study provides the first evidence that the decline of plant diversity following nutrient enrichment could occur simply if *r* is jeopardized in the first place.Specifically, N addition and soil acidification decreased *r* dramatically, and both had much stronger effects when NH_4_^+^ rather than NO_3_^-^ was applied. More importantly, the adverse effects of both on *r* were stronger for legumes than forbs and grasses. This finding is generally in line with the increase of a few dominant grasses (and forbs) at the expense of all legumes (and most grasses and forbs) with addition of N (especially NH_4_^+^ decreasing soil pH considerably, compared to NO_3_^-^) in the Park Grass (39) and many other N-addition experiments (2, 5). Conversely, *r* of some species (especially legumes) rebounded after N addition was withheld, causing plant diversity to recover (40). However, the recovery was greater for N*2PKNaMg (NO_3_^-^) than N2PKNaMg (NH_4_^+^) where soil pH remained low. Moreover, P addition increased *r* of legumes more than forbs and grasses. This is in line with the finding that P addition (especially together with K) had minor effects on plant diversity but increased legumes in the Park Grass (39) and also other fertilization experiments (41).

The responses of *r* varied among different functional groups probably for the following reasons. First, legumes are N-sensitive but P-prone, and N fixation decreases with N addition but decreases with P addition (41). Second, the smaller species (most legumes and forbs and some grasses) experience lower growth but higher mortality under the shadow of a few increasingly taller grasses due to the intensified light asymmetry after N addition (19). Third, legumes are much more sensitive to lower soil pH and metal toxicity (e.g., Al^3+^, Mn^2+^) thereof than grasses and forbs (2, 42). Thus, *r* of most species declined with nutrient addition and soil acidification but the extent differed among functional groups. Accordingly, *r* effectively explained the changes in species richness following the various treatments, which provides novel quantitative evidence for both competitive asymmetry and soil acidification hypotheses.

### Coexistence under nutrient enrichment

Coexistence hinges on the balance between niche differences and competitive asymmetries (or differences). In theory, nutrient enrichment could disrupt the balance to cause species loss by either decreasing niche differences or intensifying competitive asymmetries. The niche dimension hypothesis stresses the former (7), but it has been questioned whether the “hypothetical” niche dimensions (sensu 13) could indeed be reduced with nutrient addition (6, 18). Our study clearly demonstrated that niche differences did not decrease and even tended to increase with nutrient addition, so did structural niche differences. Moreover, niche differences increased with soil acidification. The additional analyses also showed that the effect of the number of added nutrients (niche dimensions, *sensu* Harpole & Tilman 2007) on niche differences was non-significant (χ^2^=0.001, p=0.970). Accordingly, altered niche differences poorly explained species richness.

Interestingly, we found that per-capita intra- and inter-specific competition (*a*_*ii*_ and *a*_*ij*_) both decreased dramatically with nutrient addition, suggesting that belowground competition becomes much less intense when soil nutrients are more abundant (8, 43). Besides, *a*_*ij*_ decreased more than *a*_*ii*_, and this is why niche differences 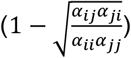 tended to increase with nutrient addition. Hence, we speculate that nutrient addition could not change “limiting” resources into “non-limiting” ones to remove niche axes. More essentially, this casts some doubts on the application of the *Law of the Minimum* (44) (in which the niche dimension hypothesis is ultimately rooted) to define how plants compete for abiotic resources (45, 46). In addition, soil acidification increased *a*_*ii*_ and *a*_*ij*_ possibly because it can make nutrients less available and impair species’ ability to utilize nutrients (26). Lastly, *a*_*ii*_ and *a*_*ij*_ also decreased with community biomass (productivity) but the effects became negligible when including individual nutrients as covariates (Fig. 6). This indicates that, not as often hypothesized (47), biomass is a poor indicator of competition intensity under nutrient enrichment.

**Figure 6.**
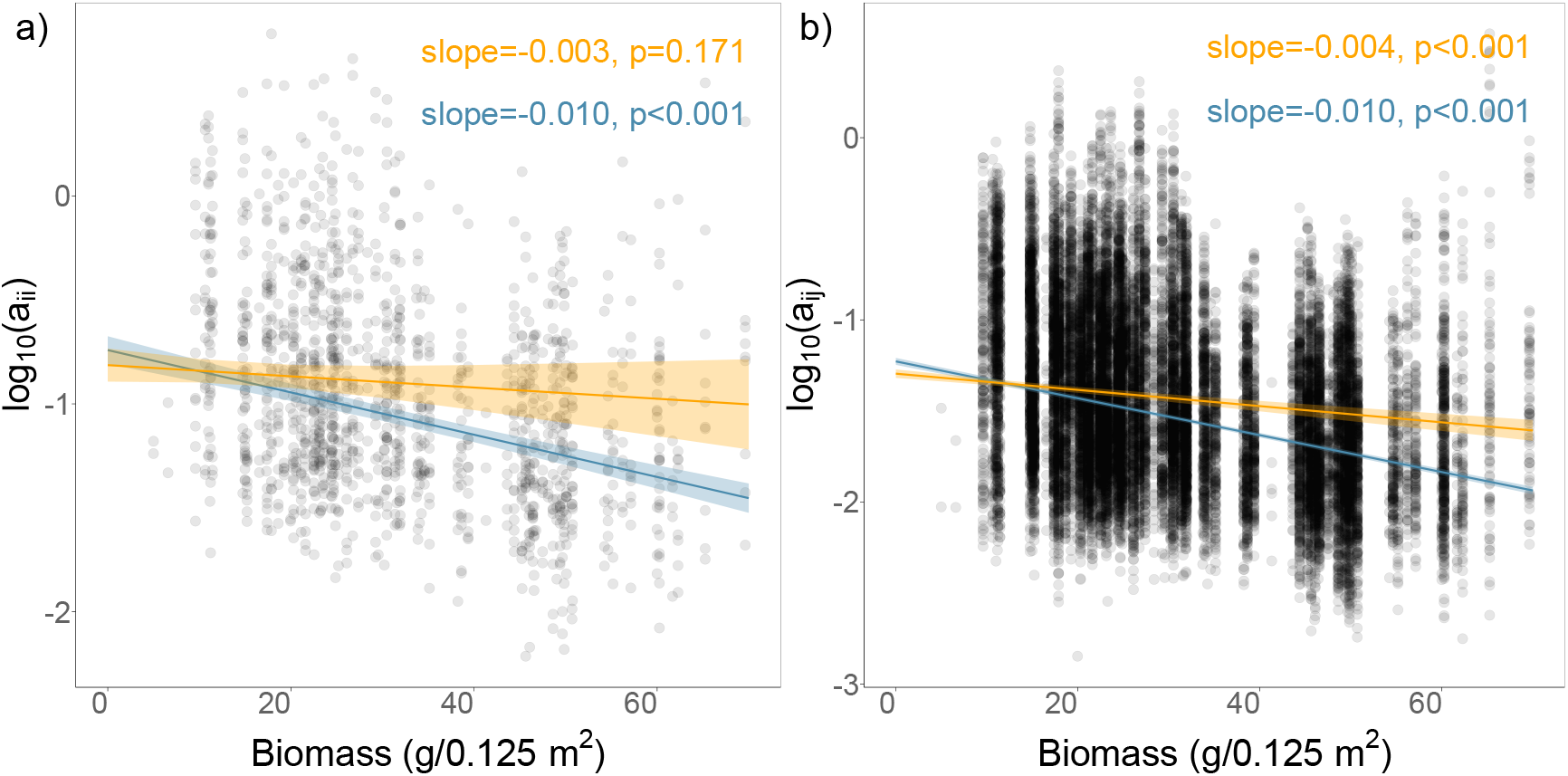
The effects of community biomass on per-capita intraspecific and interspecific competition (a: *a*_*ii*_ ; b: *a*_*ij*_). The blue lines with errors are predicted from the linear models in which the explanatory variables are community biomass (of each plot), soil pH and functional group, while the orange lines with errors are predicted from the linear models in which all individual nutrients (i.e., N type, P, K, NaMg) are included as additional covariates.

Nutrient enrichment could also cause species loss by intensifying competitive asymmetries. Despite the empirical evidence that nutrient enrichment could intensify the asymmetry of light competition (8, 19, 48) and belowground competition for soil nutrients (9, 24), little known is how this plays out to decrease plant diversity through population dynamics and species interactions. By fitting dynamic competition models with time-series data, we found that both nutrient addition and soil acidification greatly intensified competitive asymmetries among different functional groups. Briefly, N or P addition and soil acidification made grasses more competitive than forbs and legumes (and also forbs more competitive than legumes), while K or NaMg addition made legumes more competitive than forbs and grasses. Clearly, the findings are in line with the abundance changes of the three functional groups generally observed in many fertilization experiments (6, 39). Thus, absolute competitive differences (together with *r*) effectively explained species richness, which provides additional quantitative evidence for the competitive asymmetry and soil acidification hypotheses.

The further analyses showed that competitive asymmetries or differences 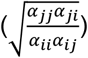 were intensified as a result of the differential changes of three functional groups in *a*_*ii*_ and *a*_*ij*_. For instance, probably because grasses are more N-limited (2) and also suffer less from the intensified light asymmetry (19) than forbs and legumes, N addition decreased the per-capita competitive effects of grasses on themselves (*a*_*ii*_) (i.e., increased carrying capacity) and on forbs or legumes (*a*_*ij*_) more than the reverse. In fact, N addition increased legumes’ *a*_*ii*_ (i.e., decreased carrying capacity) and also their *a*_*ij*_ on grasses or forbs. However, as the extent that carrying capacity (i.e., equilibrial biomass in monoculture) increased more for grasses than forbs and legumes overrode the extent that *a*_*ij*_ of grasses on forbs and legumes decreased more than the reverse, N addition made the overall competitive effects (biomass × per-capita effects) on forbs and legumes greater than those on grasses, i.e., grasses more competitive than forbs and legumes. For the reasons alike, N addition made forbs more competitive than legumes and K addition made legumes more competitive than forbs and grasses. Moreover, as grasses are more tolerant of lower soil pH than forbs and legumes (2, 39), soil acidification made grasses more competitive than forbs and legumes by increasing grasses’ *a*_*ii*_ (i.e., decreasing carrying capacity) and also their *a*_*ij*_ on forbs and legumes less than the reverse. For the similar reasons, soil acidification also made forbs more competitive than legumes. Given all this, future research should further explore the aboveground and belowground biological bases of how emergent population-level competitive asymmetries are intensified under nutrient enrichment.

## Conclusion

An unresolved fundamental mystery in ecology is what causes the general loss of plant diversity following nutrient enrichment. Ecologists have long applied classical coexistence theory to wrestle with this conundrum, but the proposed hypotheses remain highly controversial. Our study for the first time made use of modern coexistence theory and its theoretical linkages to the essential elements of the three main hypotheses to unravel the complexity with the famous long-term Park Grass. Overall, our quantitative findings strongly favored the competitive asymmetry and soil acidification hypotheses over the niche dimension hypothesis. Moreover, our study explicitly demonstrated that *existence* and *coexistence* both played a critical role in the process of species loss under nutrient enrichment. All this provides rigorous and quantitative explanations for the general findings in numerous nutrient-addition experiments (see a new meta-analysis, Band *et al*. 2022). In total, our study advances the knowledge of the fundamental mechanisms driving the response of plant diversity to nutrient deposition in nature. Finally, we believe that our study will inspire future studies to apply modern coexistence theory to more quantitatively understand, predict and manage the responses of plant communities to other global change factors (e.g., climate change, land use change) in nature.

## Materials and Methods

### Park Grass

The Park Grass Experiment is the world’s longest-running fertilization experiment, established in 1856 on a speciose mesotrophic meadow of c. 2.8 ha located at Harpenden, England (51°48’ N, 0°22’ W; elevation: 133 m; MAT: 9.34 °C, MAP: 709 mm) (49, 50). In the experiment, there exist a total of about 64 species from 19 families, which belong to three functional groups: 17 grasses, 42 forbs and 5 legumes (Table S1). The Park Grass is an ideal platform to address our research problems as it has various treatments of nutrient addition (CK and 14 different fertilization treatments by selectively combining N as NH_4_^+^ or NO_3_^-^ each with three levels, P, K, Na, Mg and Si) fully crossed with four levels of soil pH adjusted by liming (unlimed, 5, 6 and 7) (51) (Box S1). Over 150 years, the various treatments had caused dramatic changes in the diversity and composition of plant communities (Table S2 and Fig. S1) (39). For instance, N addition caused the decline of plant diversity with some grasses or forbs increased at the expense of all legumes and most grasses and forbs, especially when adding NH_4_^+^ (causing severe soil acidification) rather than NO_3_^-^. Increasing soil pH by liming partially recovered plant diversity. Moreover, addition of P (particularly together with K) marginally affected plant diversity but significantly increased the abundance of legumes. Lastly, the experiment has a unique “transient” treatment in which N addition was withheld in half of N2PKNaMg (NH_4_^+^) and N*2PKNaMg (NO_3_^-^) since 1990, which has made community diversity and composition bounce back (40). Species-level biomasses have been surveyed > 30 times since 1856 (52, 53), and the time series can be used to fit the Lotka-Volterra competition models (see below). For more details about the Park Grass, see SI Appendix S1.

### Theory

According to modern coexistence theory, niche and competitive differences can be defined with the Lotka-Volterra competition model (Eqn. 1)

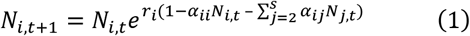

where *N*_*i,t*+1_ and *N*_*i,t*_ are biomasses of species *i* at time *t* + 1 and *t* respectively and *N*_*j,t*_ is biomass of species *j* at time *t, s* is total number of species in the system, and *r*_*i*_ is intrinsic (or maximum) population growth rate of species *i* (in the absence of intra- and inter-specific competition). Moreover, *α*_*ii*_ is intraspecific competition coefficient measuring the per-capita competitive effect of species *i* on itself and *α*_*ij*_ is interspecific competition coefficient measuring the per-capita competitive effect of species *j* on species *i*.

Niche difference between species *i* and *j* is calculated as 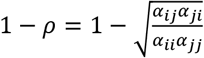 (where *ρ* is niche overlap), and it essentially measures the strength of intraspecific relative to interspecific competition. In contrast, competitive difference is calculated as 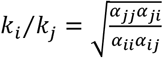, which quantifies the overall difference of competitive ability between species *i* and *j*. For example, *k*_*i*_ *⁄k* _*j*_ > 1 signifies that species *i* is more competitive than species *j*, as the intra- and inter-specific competitive effects on species *j* (*α*_*jj*_*α*_*ji*_) are overall greater than on species *i* (*α*_*ii*_ *α*_*ij*_). It is also essential to note that *k*_*i*_*⁄k*_*j*_ implicitly reflects the biomass difference of the two competing species at equilibrium, i.e., carrying capacity (*K*: *K*_*i*_ = 1/*α*_*ii*_, *K*_*j*_ = 1/*α*_*jj*_) constrained by interspecific competition (*α*_*ij*_, *α*_*ji*_). For instance, if species *i* has a greater carrying capacity than species *j* (i.e., *α*_*ii*_ < *α*_*jj*_) and exerts a stronger per-capita competitive effect on species *j* than the reverse (*α*_*ji*_ > *α*_*ij*_), species *i* will have greater biomass (i.e., “appear” to be more competitive from an empirical perspective) than species *j*. Note that, compared to niche difference that is *directionless*, competitive difference is *directional* as it becomes the reciprocal if species *i* and *j* exchange. Species *i* and *j* are competitively symmetric if *k*_*i*_*⁄k*_*j*_ = 1. However, if *k*_*i*_*⁄k*_*j*_ deviates from 1 (i.e., 1 → 0 or 1 → +∞), species *i* and *j* become competitively more asymmetric, i.e., one species becomes more competitive than the other. In theory, 1) species *i* and *j* stably coexist when niche difference (or overlap) contains competitive difference, i.e., 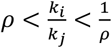 ;2) one species competitively excludes the other if 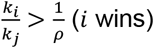 or 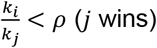;3)priority effects occur if 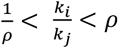, where one species with greater initial biomass wins (54). Lastly, analogous to the pairwise ones, multispecies structural niche and competitive differences were also proposed (55).

### Model fitting

The fitting of all the models was executed using Bayesian inference with the “rstan” package (56) in R 4.1.2 (57), and see SI Appendix S2 for the details. Briefly, to make the best use of the rich long-term data to achieve our research goals, we organized all the time-series data into the “recent” and “longul” subsets. The “recent” included all the plots from 1965 to 2000 during which the four levels of soil pH were applied in each treatment of nutrient addition, while the “longul” is a smaller but longer dataset with “unlimed” vs “limed” plots from 1856 to 2000. This organization allowed us to best explore the independent effects of nutrient addition and soil acidification and their interactions while still capitalizing on the long-term merit of the Park Grass. To further facilitate the models’ fitting, we merged the species that are rare in a plot as “rare species”, which substantially reduced the number of explicitly fitted species per plot without affecting model performance.

Separately for each subset, we then fitted ln-transformed Lotka-Volterra competition models (Eqn. 2) with time-series data of each plot.

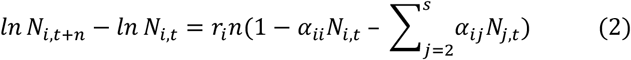

where *n* is the number of years between two consecutive time points, and other parameters are the same as in Eqn. 1. Specifically, we fitted linear mixed models in which we set *ln N*_*i,t*+*n*_ − *ln N*_*i,t*_ (i.e., the difference in ln-transformed biomasses of species *i* in two time points) as the dependent variable, *N*_*i,t*_ and *N*_*j,t*_ as the independent variables, and “period” (from *t* to *t* + *n* is a period) as the random variable. We also tried including climatic variables (precipitation, temperature) as additional random variables but the results were almost the same (not shown), so we left them out for simplicity. This indicates that the “period” has virtually captured the effects of climatic fluctuations. Thus, our models in fact allow us to explore fluctuation-independent coexistence mechanisms (i.e., niche and competitive differences) in which we are interested while controlling for fluctuation-dependent ones (e.g., temporal storage effects) (27).

The diagnostics of the model fits (*n*_*eff*_, 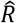 and their SDs) suggest that the fitted models are overall reliable (Table S3). In addition, we applied the “leave-one-year-out” cross-validation to assess the goodness-of-fit following Tredennick, Hooten and Adler (58). The average high accuracy (*ρ*) (recent: 0.50; longul: 0.60) and low error (recent: 1.70; longul: 0.88; unit: g/0.125 m^2^) of model predictions further corroborate the adequate reliability of the fitted models (Fig. S4). Lastly, we derived all the variables required for further analyses by taking the means of posteriors of respective parameters (medians gave identical results). In total, we obtained 36 explicitly fitted species and 11455 species pairs across all plots and treatments (CK: 2883; fertilized: 8572). Note that we removed a small number of pairs to ensure that all the included pairs occur in at least two treatments to avoid the potential confounding effects of community composition across treatments in the further analyses.

### Statistical analyses

First, we used linear models to analyze how different treatments and their interactions affected intrinsic population growth rate (*r*), niche difference 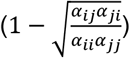, competitive difference 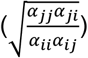, and per-capita intraspecific and interspecific competition (*a* _*ii*_ and *a* _*ij*_). All the response variables, except for *r* (square root transformed) and niche difference, were log_10_-transformed to meet the assumptions for residuals. As the explanatory variables, we included N type (0, NH_4_^+^, NO_3_^-^), P (0, 1), K (0, 1), NaMg (0, 1), soil pH, “functional group” and all possible interactions. We included NaMg as a whole because Na and Mg were always added simultaneously, and we dropped Si because its effects were all non-significant. Moreover, we also tried including N amount (three levels) but the results were very similar to those of N type (and the two were moderately confounded), so we decided to only keep N type. The inclusion of “functional group” and its interactions with other variables allows us to explore the effect patterns at the level of functional group. For the obvious reason, however, the “functional group” variable differed for different response variables: 1) three functional groups (grass, forb and legume) for *r* and *a*_*ii*_, 2) six functional-group combinations for niche difference that is *directionless*, 3) nine functional-group permutations for both *a*_*ij*_ and competitive difference that are *directional*. Note that for *r* we run an additional model replacing N type (0, NH_4_^+^, NO_3_^-^) with N (0, 1) throughout to show how the overall effect of N addition differed for three functional groups (as the interaction for N type was negligible). For other interactions, we included all possible interactions between different nutrients (except for a few not included for logistical constraints) and all interactions between different nutrients and soil pH. For simplicity, we analyzed the merged output of the “recent” and “longul” subsets because the separate analyses of the two yielded very similar results (not shown). Moreover, to test whether withholding N addition reversed the long-term effects of N addition, we used linear models with the output from the “transient” treatment to analyze how each of the abovementioned response variables was affected by N withholding (0, 1), N type (NH_4_^+^, NO_3_^-^), soil pH, “functional group” and the interactions between N withholding and both N type and “functional group”. For all the analyses above, we also tried linear mixed-effects models including species nested within family as the random variable, but the results were almost the same (not shown). In addition to the statistical analyses, we constructed the competitive hierarchy of all species in each plot based on competitive differences to show how various treatments affected competitive hierarchies (Table S6).

Second, we used generalized linear models with Poisson distribution to analyze how plot-level species richness was explained by the changes (due to the various treatments of nutrient addition and soil pH) in plot-level *r* (averaged across all species), niche difference and absolute competitive difference (i.e., |competitive difference|) (averaged across all species pairs), and their interactions. We included absolute competitive difference because it reflects absolute competitive asymmetry and could explain species richness more straightforwardly than competitive difference that is directional. For all the models, we used the log-likelihood-ratio tests to assess the significance of explanatory variables and their interactions. The detailed results of the statistical models are provided in SI Appendix S3. We illustrated the significant and marginally significant main and interaction effects with the “sjPlot” and “ggplot2” R packages (59, 60). The analyses and results of structural niche and competitive differences are provided in SI Appendix S4. All the statistical analyses were executed in R 4.1.2 (57).

## Acknowledgments

This work was supported by the National Natural Science Foundation of China (No. 31971433), the Strategic Priority Research Program of the Chinese Academy of Sciences (No. XDA26020202) and the Fundamental Research Funds for the Central Universities (No. lzujbky-2021-ey02 & lzujbky-2019-32). We are grateful to the Rothamsted Research and the electronic Rothamsted Archive (e-RA) for sharing the valuable data of the Park Grass and Sarah Perryman for helping arrange the data. We thank Xiang Liu and Shaopeng Li for helpful discussions and comments on earlier versions of the manuscript.

