## Supporting Information for "Species existence and coexistence under nutrient enrichment in the Park Grass"

* Yanhao Feng

**This PDF file includes:**

Supplementary text
Figures S1 to S7
Tables S1 to S8
Box S1
SI References

**Appendix S1 Detailed information about the Park Grass**

*Species and treatments*

The Park Grass Experiment was established on a species-rich meadow that had been a permanent pasture for at least 100 years (1). Across all plots and years, there exist a total of about 60 species that belong to roughly 20 different families and can be categorized as three functional groups, i.e. grass, forb and legume (2) (Table S1). Almost all species are perennials, with a few exceptions of annuals and biennials (*Crepis capillaris*, *Linum catharticum*). The overall relative abundance of the species ranges from > 10% for a few dominant grasses (*Agrostis capillaris*, *Arrhenatherum elatius, Festuca rubra*) through 1-10% for more than a dozen common or relatively common species (e.g. *Anthoxanthum odoratum*, *Leontodon hispidus*, *Lathyrus pratensis*) to <1% for many rare forbs and legumes (e.g. *Leontodon autumnalis*, *Carex caryophyllea*, *Sanguisorba minor*, *Lotus corniculatus*) (Table S1). So, the average number of species at the level of plots without nutrient addition (CK) is about 40, and roughly 20 species among them already comprise > 95% of total biomasses across plots (Table S2).

As shown in Box S1, the experiment consists of 15 different treatments of nutrient addition including CK (i.e. "Nil") and 14 fertilization treatments by selectively combining N as $(NH_4)_2SO_4$ ($NH_4^+$) or $NaNO_3$ ($NO_3^-$) (each with three levels), P ($CaP_2H_4O_8$), K ($K_2SO_4$), Na ($Na_2SO_4$), Mg ($MgSO_4$) and Si ($Na_2SiO_3$), with the number of added nutrients across treatments ranging from 0 (CK) to 6 (N3PKNaMgSi). There is one plot per treatment, except for CK with three replicates (plot 2/2, 3 and 12) and PKNaMg with two replicates (plot 7 and 15), totalling 19 original plots (75 to 634 $m^2$). The size of the plots, in addition to the reasonable uniformity of the meadow prior to the experiment (1), compensates for the deficiency in the experimental design from the perspective of modern statistics (e.g. lack of replication and randomization) (3). Most of the treatments started before the 1880s, with a few exceptions that began after 1900 (i.e. N2KNaMg, N1PKNaMg, K). Note we left out K (1996-, plot 2/1) from our analyses as the data are insufficient to fit models. Moreover, note N addition was withheld since 1990 in half of plots with two treatments (plots 9 and 14), i.e. N2PKNaMg and N*2PKNaMg with N added as $(NH_4)_2SO_4$ and $NaNO_3$ respectively. These unique treatments signified as (N2)PKNaMg (plot 9/1) and (N*2)PKNaMg (plot 14/1) were introduced to test whether withholding N could make community diversity and composition recover.

The soil is clay silty loam classified as mildly acidic Luvisol with poor nutrient status. The soil pH (0-23cm) was about 5.5 before the experiment starts, and now it becomes c. 5.0 in the unlimed "d" quarter-plots of CK (i.e. Nil) treatments, and nutrient addition further reduces soil pH especially when N is added as $(NH_4)_2SO_4$ (Table S2). To dilute soil acidity, a small amount of lime was added into each plot in the late 1880s. Since 1903, each plot was equally divided into two half-plots, one applied with lime (4t $CaCO_3$ $ha^{-1}$) every four years, the other as the unlimed control. From 1965, each half-plot was further divided into two quarter-plots (limed: "a" and "b"; unlimed: "c" and "d"), with soil pH of the "a", "b", and "c" maintained at 7, 6, and 5 respectively by liming every three years, and with the "d" unlimed (4). This design allows the exploration of the effects of nutrient addition independent of soil acidification. For simplicity, hereafter we use plot to substitute for "plot", "half-plot" and "quarter-plot", unless otherwise specified. More detailed information about the Park Grass Experiment can be found in the electronic Rothamsted Archive (www.era.rothamsted.ac.uk/Park).

*Community diversity and composition*

The community composition of each plot has been surveyed in early June on more than 30 occasions since the experiment began. From 1856 to 1990, species-level biomass was determined in selected years (> 20 occasions in total) by taking several small samples (later merged) at regular intervals along a transect in each plot. From 1991 to 2000, species-level biomass was determined annually by taking six samples, each with a quadrat (50cm × 25cm) randomly placed in each plot (2).

Total biomass (hay yield) of each plot was also harvested in early June every year since 1856, although the harvesting methods changed with agricultural technology development (5). The original purpose of the Park Grass Experiment was to investigate the effects of fertilization by inorganic nutrients (or organic manures) on improving the hay yield (see total biomass in Table S2) (1). However, a more interesting phenomenon quickly emerged within just a few years that fertilization had dramatic effects on the community diversity and composition of the meadow that had been reasonably uniform for a very long term. Later analyses suggest that this is not only because of nutrient addition per se, but also because of soil acidification concomitant with nutrient addition (as mentioned above, this is why soil pH treatments were additionally introduced) (3).

Over 150 years, the various treatments of nutrient addition crossed with soil pH adjustments have produced a wide variety of plant communities with contrasting diversity and composition (Table S2 and Fig. S1). For example, the CK plots (plots 3, 2/2 and 12) still contain 38-45 species and roughly maintain the original composition, but there only remain a couple of grasses (e.g. *A. capillaris*, *Holcus lanatus*) in the unlimed “d” plots of treatments such as N3PKNaMgSi (plot 11/2). In general, addition of different nutrients has distinct effects, and their effects appear to interact with soil pH treatments that alone regulate the rise and decline of acid resistant species such as some grasses (e.g. *A. odoratum*) and acid sensitive species including all legumes and most forbs (Table S2 and Fig. S1). The addition of N overall causes the dramatic decline of species richness and change in community composition with some grasses or forbs increased at the expense of most grasses and forbs and all legumes (see all plots where N is added in Table S2 and Fig. S1), but the effects of $(NH_4)_2SO_4$ ($NH_4^+$) are much greater than those of $NaNO_3$ ($NO_3^-$). This is mainly because soil pH drops to as low as 3.6 when $(NH4)_2SO_4$ is applied (e.g. plot 10) but almost has no change when $NaNO_3$ is applied (e.g. plot 16). As a result, in most plots where $(NH4)_2SO_4$ is applied, almost all legumes and forbs vanish and only a few grasses (e.g. *A. odoratum* and *A. capillaris*) become dominant as they are able to greatly tolerate the toxicity of $Al^{3+}$ whose concentration increases at low soil pH (Table S2 and Fig. S1) (6). Following this, increasing soil pH by liming has much greater effects for the plots with $(NH4)_2SO_4$ (e.g. plot 9/2) than those with $NaNO_3$ (e.g. plot 14/2). However, the liming only leads to the recovery of some species (e.g. some forbs) but not others (e.g. legumes), indicating the adverse effects of N addition independently of soil acidification especially on legumes. The addition of P alone (plot 4/1) and PNaMg (plot 8) only has minor effects on species richness and community composition, but it tended to increase the abundance of three legumes (*L. pratensis*, *Trifolium pratense* and *Trifolium repens*), especially when it is added together with K (plot 7) (Table S2). In line with the latter, it is clear that the combination of more different nutrients overall yields much stronger effects (Fig. S1). Moreover, the community diversity and composition bounced back, especially legumes increased, when N was withheld in the (N2)PKNaMg (plot 9/1) and (N*2)PKNaMg (plot 14/1) and nitrogen deposition naturally declined (7). Finally, interestingly small changes have also occurred for the CK plots without fertilization over 150 years, with some grasses (*Lolium perenne*, *Holcus lanatus*) gradually replaced by other grasses (*A. capillaris*, *F. rubra*) and with some forbs (e.g. *L. hispidus*, *Centaurea nigra*) increased over time, possibly due to the dynamics of nutrient deposition in nature.

**Table S1** Species of the Park Grass Experiment. The 64 species are from 19 families that belong to three functional groups: grass, forb and legume. The relative abundance (RA, %) of each species is averaged from 1991 to 2000 across all plots.

| ID | Group | Family | Code | Species | Common name | Growth form | RA |
|---|---|---|---|---|---|---|---|
| 1 | grass | Poaceae | *Agr.cap* | *Agrostis capillaris* | common bent | perennial | 12.82 |
| 2 | grass | Poaceae | *Alo.pra* | *Alopecurus pratensis* | meadow foxtail | perennial | 8.96 |
| 3 | grass | Poaceae | *Ant.odo* | *Anthoxanthum odoratum* | sweet vernal grass | perennial | 5.47 |
| 4 | grass | Poaceae | *Arr.ela* | *Arrhenatherum elatius* | tall oat grass | perennial | 11.91 |
| 5 | grass | Poaceae | *Bri.med* | *Briza media* | quaking grass | perennial | 0.41 |
| 6 | grass | Poaceae | *Bro.hor* | *Bromus hordeaceus* | soft brome | perennial | 0.27 |
| 7 | grass | Poaceae | *Cyn.cri* | *Cynosurus cristatus* | crested dog's-tail | perennial | <0.01 |
| 8 | grass | Poaceae | *Dac.glo* | *Dactylis glomerata* | orchard grass | perennial | 4.12 |
| 9 | grass | Poaceae | *Des.ces* | *Deschampsia cespitosa* | tufted hair grass | perennial | <0.01 |
| 10 | grass | Poaceae | *Fes.pra* | *Festuca pratensis* | meadow fescue | perennial | 0.10 |
| 11 | grass | Poaceae | *Fes.rub* | *Festuca rubra* | red fescue | perennial | 12.44 |
| 12 | grass | Poaceae | *Hel.pub* | *Helictotrichon pubescens* | downy oat-grass | perennial | 0.97 |
| 13 | grass | Poaceae | *Hol.lan* | *Holcus lanatus* | velvet grass | perennial | 9.08 |
| 14 | grass | Poaceae | *Lol.per* | *Lolium perenne* | perennial ryegrass | perennial | 0.50 |
| 15 | grass | Poaceae | *Poa.pra* | *Poa pratensis* | Kentucky bluegrass | perennial | 1.29 |
| 16 | grass | Poaceae | *Poa.tri* | *Poa trivialis* | rough bluegrass | perennial | 1.65 |
| 17 | grass | Poaceae | *Tri.fla* | *Trisetum flavescens* | yellow oatgrass | perennial | 0.23 |
| 18 | forb | Apiaceae | *Ant.syl* | *Anthriscus sylvestris* | cow parsley | perennial | 1.71 |
| 19 | forb | Apiaceae | *Con.maj* | *Conopodium majus* | pignut | perennial | 0.41 |
| 20 | forb | Apiaceae | *Her.sph* | *Heracleum sphondylium* | hogweed | perennial | 2.38 |
| 21 | forb | Apiaceae | *Pim.sax* | *Pimpinella saxifraga* | burnet saxifrage | perennial | 0.08 |
| 22 | forb | Asteraceae | *Ach.mil* | *Achillea millefolium* | common yarrow | perennial | 1.07 |
| 23 | forb | Asteraceae | *Bel.per* | *Bellis perennis* | common daisy | perennial | <0.01 |
| 24 | forb | Asteraceae | *Cen.nig* | *Centaurea nigra* | lesser knapweed | perennial | 2.84 |
| 25 | forb | Asteraceae | *Cre.cap* | *Crepis capillaris* | smooth hawksbeard | annual or biennial | 0.02 |
| 26 | forb | Asteraceae | *Hyp.rad* | *Hypochoeris radicata* | cat's ear | perennial | 0.02 |
| 27 | forb | Asteraceae | *Leo.aut* | *Leontodon autumnalis* | fall dandelion | perennial | 0.07 |
| 28 | forb | Asteraceae | *Leo.his* | *Leontodon hispidus* | bristly hawkbit | perennial | 3.36 |
| 29 | forb | Asteraceae | *Pil.off* | *Pilosella officinarum* | mouse-ear hawkweed | perennial | 0.02 |
| 30 | forb | Asteraceae | *Sen.jac* | *Senecio jacobaea* | common ragwort | perennial | <0.01 |
| 31 | forb | Asteraceae | *Tar.off* | *Taraxacum officinale* | common dandelion | perennial | 1.58 |

| | | | | | | | |
|---|---|---|---|---|---|---|---|
| 32 | forb | Asteraceae | *Tra.pra* | *Tragopogon pratensis* | meadow goat's-beard | biennial | 0.66 |
| 33 | forb | Caprifoliaceae | *Kna.arv* | *Knautia arvensis* | field scabious | perennial | 0.18 |
| 34 | forb | Caryophyllaceae | *Cer.fon* | *Cerastium fontanum* | mouse-ear chickweed | perennial | 0.09 |
| 35 | forb | Caryophyllaceae | *Ste.gra* | *Stellaria graminea* | common starwort | perennial | <0.01 |
| 36 | forb | Caryophyllaceae | *Ste.med* | *Stellaria media* | common chickweed | annual or perennial | <0.01 |
| 37 | forb | Cyperaceae | *Car.car* | *Carex caryophyllea* | vernal sedge | perennial | 0.07 |
| 38 | forb | Cyperaceae | *Car.fla* | *Carex flacca* | blue-green sedge | perennial | 0.03 |
| 39 | forb | Juncaceae | *Luz.cam* | *Luzula campestris* | field wood-rush | perennial | 0.25 |
| 40 | forb | Lamiaceae | *Aju.rep* | *Ajuga reptans* | blue bugle | perennial | 0.02 |
| 41 | forb | Lamiaceae | *Pru.vul* | *Prunella vulgaris* | common self-heal | perennial | <0.01 |
| 42 | forb | Lamiaceae | *Sta.off* | *Stachys officinalis* | common hedgenettle | perennial | 0.02 |
| 43 | forb | Liliaceae | *Fri.mel* | *Fritillaria meleagris* | snake's head | perennial | <0.01 |
| 44 | forb | Linaceae | *Lin.cat* | *Linum catharticum* | purging flax | annual | 0.02 |
| 45 | forb | Ophioglossaceae | *Oph.vul* | *Ophioglossum vulgatum* | adder's-tongue | perennial | <0.01 |
| 46 | forb | Plantaginaceae | *Ver.cha* | *Veronica chamaedrys* | germander speedwell | perennial | 0.01 |
| 47 | forb | Plantaginaceae | *Pla.lan* | *Plantago lanceolata* | ribwort plantain | perennial | 4.01 |
| 48 | forb | Polygonaceae | *Rum.ace* | *Rumex acetosa* | common sorrel | perennial | 1.80 |
| 49 | forb | Primulaceae | *Pri.ver* | *Primula veris* | common cowslip | perennial | <0.01 |
| 50 | forb | Ranunculaceae | *Ran.acr* | *Ranunculus acris* | meadow buttercup | perennial | 1.94 |
| 51 | forb | Ranunculaceae | *Ran.aur* | *Ranunculus auricomus* | goldilocks buttercup | perennial | 0.02 |
| 52 | forb | Ranunculaceae | *Ran.bul* | *Ranunculus bulbosus* | bulbous buttercup | perennial | 0.04 |
| 53 | forb | Ranunculaceae | *Ran.fic* | *Ranunculus ficaria* | pilewort | perennial | <0.01 |
| 54 | forb | Rosaceae | *Agr.eup* | *Agrimonia eupatoria* | common agrimony | perennial | 0.01 |
| 55 | forb | Rosaceae | *Fil.ulm* | *Filipendula ulmaria* | meadowsweet | perennial | <0.01 |
| 56 | forb | Rosaceae | *Pot.rep* | *Potentilla reptans* | creeping cinquefoil | perennial | <0.01 |
| 57 | forb | Rosaceae | *Pot.ste* | *Potentilla sterilis* | strawberryleaf cinquefoil | perennial | <0.01 |
| 58 | forb | Rosaceae | *San.min* | *Sanguisorba minor* | salad burnet | perennial | 0.28 |
| 59 | forb | Rubiaceae | *Gal.ver* | *Galium verum* | yellow spring bedstraw | perennial | 0.03 |
| 60 | legume | Fabaceae | *Lat.pra* | *Lathyrus pratensis* | meadow vetchling | perennial | 2.19 |
| 61 | legume | Fabaceae | *Lot.cor* | *Lotus corniculatus* | common bird's-foot trefoil | perennial | 0.37 |
| 62 | legume | Fabaceae | *Ono.rep* | *Ononis repens* | common restharrow | perennial | 0.02 |
| 63 | legume | Fabaceae | *Tri.pra* | *Trifolium pratense* | red clover | perennial | 4.09 |
| 64 | legume | Fabaceae | *Tri.rep* | *Trifolium repens* | white clover | perennial | 0.06 |

**Box S1** Treatments of nutrient addition (left) and plot layout of Park Grass Experiment (right). No., plot and year indicate number of added nutrients, plot code and starting time of the 15 different treatments respectively. Previous treatments of them (if any) are also indicated.

| No. | ID | Treatment | Plot | Year | Previous treatments and comments |
|---|---|---|---|---|---|
| 0 | 1 | CK (i.e. Nil) | 2/2 | 1864- | FYM (1856-1863) |
| | 1 | CK (i.e. Nil) | 3 | 1856- | |
| | 1 | CK (i.e. Nil) | 12 | 1856- | |
| 1 | 2 | N1 | 1 | 1856- | with FYM (1856-1863) |
| | 3 | N*1 | 17 | 1858- | |
| | 4 | P | 4/1 | 1856- | |
| 2 | 5 | N2P | 4/2 | 1856- | |
| 3 | 6 | PNaMg | 8 | 1862- | PKNaMg (1856-1861) |
| 4 | 7 | PKNaMg | 7 | 1856- | |
| | 7 | PKNaMg | 15 | 1876- | N*2 (1858-1875) |
| | 8 | N2PNaMg | 10 | 1856- | with K (1856-1861) |
| | 9 | N2KNaMg | 18 | 1905- | |
| 5 | 10 | N1PKNaMg | 6 | 1972- | N2 (1856-1868), PKNaMg (1869-1964) |
| | 11 | N*1PKNaMg | 16 | 1858- | no P (1866, 1867) |
| | 12 | N2PKNaMg | 9/2 | 1856- | |
| | 12 | (N2)PKNaMg | 9/1 | 1990- | i.e. withholding N2 since 1990 |
| | 13 | N*2PKNaMg | 14/2 | 1858- | |
| | 13 | (N*2)PKNaMg | 14/1 | 1990- | i.e. withholding N*2 since 1990 |
| | 14 | N3PKNaMg | 11/1 | 1882- | N4PKNaMg (1856-1858), N2PKNaMg (1859-61), N4PKNaMg (1862-1881) |
| 6 | 15 | N3PKNaMgSi | 11/2 | 1882- | N4PKNaMg (1856-1858), N2PKNaMg (1859-61), N4PKNaMgSi (1862-1881) |

**Treatments ($ha^{-1}$ $year^{-1}$, all applied in spring, except for N and N* applied in winter)**

**N1, N2 and N3:** 48, 96 and 144 kg N as $(NH_4)_2SO_4$

**N*1, N*2 and N*3:** 48, 96 and 144 kg N as $NaNO_3$

**P:** 35 kg P as $CaP_2H_4O_8$

**K:** 225 kg K as $K_2SO_4$

**Na:** 15 kg Na as $Na_2SO_4$

**Mg:** 10 kg Mg as $MgSO_4$**Si:** 450 kg of $Na_2SiO_3$

**FYM:** 35 t farmyard manure (every four years)

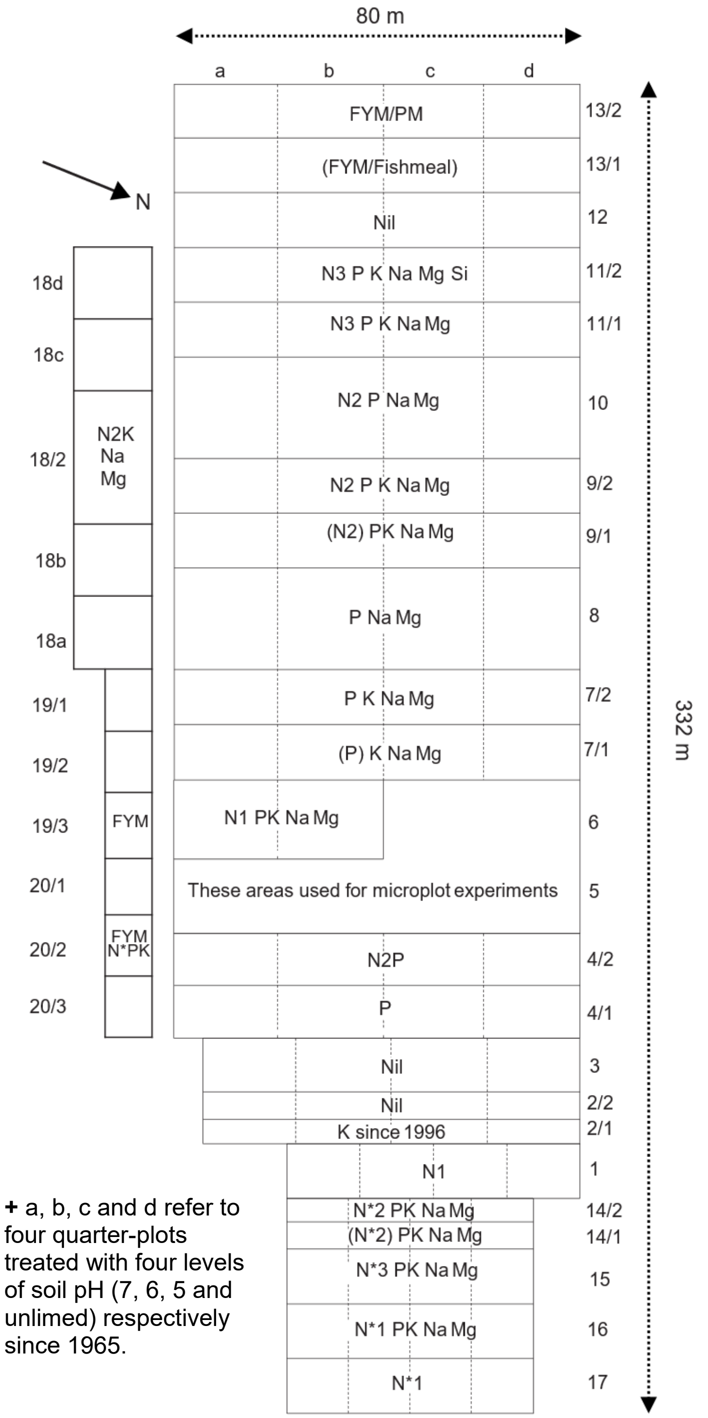


**+** a, b, c and d refer to four quarter-plots treated with four levels of soil pH (7, 6, 5 and unlimed) respectively since 1965.

**Table S2** Species richness (SR), total biomass (Bio), and relative abundance (%) of 20 representative species (for abbreviation, see Table S1), in each of the four quarter-plots (QP) within each of the plots with 15 different treatments of nutrient addition. The 20 species include eight grasses, nine forbs and three legumes, comprising > 95% of biomasses across all plots. Soil pH of the "a", "b" and "c" was maintained at 7, 6 and 5 respectively by liming every three years, with the "d" unlimed, since 1965. The SR, Bio, actual pH ($pH_a$) and relative abundance are all averaged from 1991 to 2000. Note "*" indicates RA is <1% and "-" indicates the absence of species.

| Treatment (Plot) | QP | $pH_a$ | SR | Bio | *Ac* | *Ap* | *Ao* | *Ae* | *Dg* | *Fr* | *Hl* | *Pp* | *Hs* | *Am* | *Cn* | *Lh* | *To* | *Pl* | *Ra* | *Ra* | *Sm* | *Lp* | *Lc* | *Tp* |
|---|---|---|---|---|---|---|---|---|---|---|---|---|---|---|---|---|---|---|---|---|---|---|---|---|
| | a | 6.8 | 45 | 35.0 | 12 | * | 2 | * | 3 | 15 | 2 | 1 | * | 3 | 6 | 16 | * | 8 | * | 1 | * | 1 | * | 4 |
| CK (i.e. Nil) | b | 6.2 | 43 | 38.5 | 13 | * | 3 | 1 | 4 | 16 | 2 | 1 | * | 3 | 7 | 16 | 1 | 8 | * | 2 | * | 1 | * | 2 |
| (2/2, 3, 12) | c | 5.1 | 38 | 30.0 | 30 | * | 3 | * | 1 | 34 | * | * | * | 3 | 6 | 9 | * | 3 | * | 1 | * | * | * | * |
| | d | 5.0 | 38 | 29.6 | 34 | * | 3 | * | 2 | 30 | 1 | * | * | 2 | 11 | 7 | * | 3 | * | * | * | * | * | * |
| | a | 6.7 | 35 | 36.8 | 7 | 1 | 2 | 3 | 8 | 27 | 3 | 2 | - | 1 | 8 | 8 | * | 8 | 3 | * | * | * | * | * |
| N1 | b | 5.9 | 30 | 34.0 | 24 | 1 | 4 | * | 7 | 22 | 4 | 1 | - | 2 | 12 | 6 | * | 7 | 1 | * | * | * | * | * |
| (1) | c | 4.9 | 24 | 31.3 | 35 | - | 6 | * | 1 | 47 | * | * | - | 1 | 7 | - | * | - | 1 | * | - | * | * | * |
| | d | 3.9 | 5 | 20.3 | 67 | - | 32 | - | - | * | - | - | - | - | - | - | - | - | - | - | - | - | - | - |
| | a | 6.9 | 36 | 33.7 | 13 | 1 | 1 | * | 7 | 14 | 1 | 1 | 2 | 2 | 4 | 25 | 1 | 12 | * | 1 | * | - | * | * |
| N*1 | b | 6.6 | 38 | 33.6 | 16 | 2 | 4 | * | 3 | 5 | 2 | * | 2 | 2 | 7 | 29 | * | 16 | * | 1 | * | * | * | - |
| (17) | c | 5.7 | 34 | 35.1 | 27 | 2 | 4 | - | 3 | 11 | 1 | * | 1 | 2 | 7 | 23 | 1 | 11 | 1 | 1 | * | - | - | - |
| | d | 5.7 | 36 | 34.7 | 26 | 3 | 4 | 1 | 7 | 12 | 3 | * | 1 | 1 | 10 | 9 | 1 | 8 | 1 | 3 | - | - | - | * |
| | a | 6.6 | 37 | 35.6 | 7 | 1 | 3 | 2 | 1 | 20 | 3 | 2 | * | 3 | 3 | 16 | * | 11 | 1 | 6 | * | * | * | 9 |
| P | b | 6.3 | 35 | 40.7 | 4 | * | 4 | 4 | 2 | 16 | 5 | 1 | - | 5 | 4 | 12 | * | 14 | 1 | 8 | * | * | * | 4 |
| (4/1) | c | 5.2 | 33 | 34.8 | 27 | 1 | 5 | * | 1 | 24 | 4 | * | - | 5 | 2 | 11 | * | 5 | * | 5 | * | * | * | 1 |
| | d | 5.1 | 37 | 33.1 | 27 | * | 5 | * | 2 | 22 | 5 | * | - | 3 | 3 | 14 | 1 | 5 | 1 | 5 | * | * | * | 1 |
| | a | 6.5 | 24 | 44.3 | 10 | 4 | 2 | 2 | 1 | 56 | 7 | 1 | - | 1 | * | - | 1 | 4 | 4 | 2 | * | - | - | - |
| N2P | b | 5.7 | 16 | 50.1 | 13 | 13 | 4 | 1 | * | 56 | 7 | 1 | - | - | - | * | * | - | 5 | - | - | - | - | - |
| (4/2) | c | 4.9 | 18 | 41.0 | 28 | * | 2 | - | * | 56 | 2 | 2 | - | * | - | - | * | * | 9 | - | - | - | - | - |
| | d | 3.6 | 8 | 30.3 | 34 | - | 65 | - | - | * | 1 | * | - | - | - | - | - | - | * | - | - | - | - | - |
| | a | 6.7 | 40 | 35.1 | 9 | * | 3 | 3 | 1 | 17 | 6 | 1 | * | 2 | 10 | 6 | * | 5 | 1 | 7 | * | - | * | 8 |
| PNaMg | b | 6.3 | 38 | 44.0 | 6 | * | 3 | 4 | 2 | 18 | 4 | 1 | 1 | 5 | 7 | 14 | * | 7 | 2 | 8 | * | * | * | 6 |
| (8) | c | 5.2 | 33 | 37.0 | 29 | 1 | 4 | * | 3 | 20 | 3 | * | * | 3 | 5 | 13 | * | 6 | * | 3 | * | * | * | 3 |
| | d | 5.0 | 33 | 35.6 | 30 | * | 5 | * | 1 | 20 | 7 | * | - | 4 | 4 | 9 | * | 8 | * | 4 | * | * | * | 2 |
| | a | 6.7 | 29 | 65.8 | 2 | 5 | 1 | 11 | 3 | 6 | 3 | 1 | 5 | 1 | * | - | 5 | 8 | 1 | 5 | - | 16 | - | 14 |
| PKNaMg | b | 6.1 | 31 | 59.7 | 3 | 8 | 2 | 15 | 3 | 4 | 7 | 1 | 5 | 2 | 1 | * | 3 | 10 | 1 | 5 | - | 6 | - | 15 |
| (7, 15) | c | 5.0 | 30 | 56.0 | 21 | 6 | 4 | 3 | 5 | 6 | 5 | * | * | 2 | 14 | * | 1 | 3 | 2 | 1 | - | 7 | * | 16 |
| | d | 4.8 | 28 | 44.1 | 41 | 3 | 4 | 1 | 4 | 11 | 2 | * | * | 3 | 8 | 1 | * | 4 | 1 | * | - | 3 | * | 10 |
| | a | 6.7 | 23 | 50.9 | 8 | 5 | 4 | 5 | 1 | 44 | 7 | 2 | - | * | - | - | 1 | 11 | 7 | 4 | - | - | - | - |
| N2PNaMg | b | 5.6 | 15 | 53.2 | 17 | 16 | 7 | 4 | * | 38 | 4 | 1 | - | - | - | - | * | * | 12 | * | - | - | - | - |
| (10) | c | 4.7 | 14 | 50.9 | 23 | 1 | 7 | 1 | * | 51 | 3 | 2 | - | - | - | - | - | * | 11 | * | - | - | - | 1 |
| | d | 3.6 | 3 | 43.9 | 6 | - | 80 | - | - | - | 14 | - | - | - | - | - | - | - | - | - | - | - | - | - |

| | | | | | | | | | | | | | | | | | | | | | | | | |
|---|---|---|---|---|---|---|---|---|---|---|---|---|---|---|---|---|---|---|---|---|---|---|---|---|
| | a | 6.5 | 33 | 36.5 | 14 | 1 | 1 | 6 | 11 | 14 | * | * | 3 | 2 | 11 | 11 | 3 | 8 | * | - | - | - | * | * |
| N2KNaMg | b | 6.0 | 32 | 36.0 | 30 | 3 | 1 | 1 | 6 | 15 | 1 | * | 1 | * | 25 | 4 | 1 | 5 | * | - | - | - | - | - |
| (18) | c | 4.4 | 23 | 52.5 | 27 | 3 | 3 | 1 | 12 | 16 | 7 | 1 | * | * | 7 | - | * | * | 3 | * | - | - | - | - |
| | d | 3.7 | 6 | 24.2 | 79 | - | 19 | 1 | - | * | 2 | - | - | - | - | - | - | - | * | - | - | - | - | - |
| N1PKNaMg | a | 6.6 | 28 | 59.8 | * | 8 | * | 19 | 5 | 3 | 11 | 1 | 2 | 1 | * | - | 8 | 6 | 2 | 1 | - | 8 | - | 13 |
| (6) | b | 6.0 | 27 | 56.7 | 1 | 14 | 1 | 27 | 4 | 3 | 14 | 1 | 3 | 1 | * | - | 5 | 3 | 2 | 1 | - | 7 | * | 7 |
| | a | 6.8 | 26 | 67.3 | 3 | 9 | * | 22 | 4 | 14 | 1 | 1 | 4 | 1 | - | - | 2 | 3 | 2 | 3 | - | 7 | - | 10 |
| N*1PKNaMg | b | 6.4 | 26 | 60.2 | 6 | 11 | 2 | 22 | 6 | 13 | 3 | 1 | 6 | * | - | - | 2 | 6 | 1 | 4 | - | 1 | - | 7 |
| (16) | c | 5.4 | 28 | 53.5 | 24 | 9 | 5 | 16 | 4 | 9 | 4 | * | 3 | 1 | - | - | * | 6 | * | 3 | - | 3 | - | 5 |
| | d | 5.3 | 26 | 53.1 | 33 | 8 | 4 | 14 | 3 | 5 | 4 | * | 2 | 1 | 1 | - | 1 | 6 | 1 | 2 | - | 4 | - | 6 |
| | a | 6.5 | 22 | 60.2 | 1 | 15 | * | 22 | 8 | 1 | 18 | 3 | 8 | - | - | - | 5 | * | 3 | * | - | 2 | - | * |
| N2PKNaMg | b | 5.8 | 19 | 61.0 | 1 | 26 | 2 | 30 | 6 | 3 | 15 | 3 | 5 | - | - | - | 2 | - | 2 | - | - | 2 | - | * |
| (9/2) | c | 4.8 | 17 | 49.3 | 30 | 9 | 8 | 5 | 1 | 22 | 8 | 3 | 1 | - | - | - | * | - | 2 | - | - | 5 | - | 6 |
| | d | 3.7 | 4 | 41.3 | 15 | - | 66 | - | - | * | 20 | - | - | - | - | - | - | - | - | - | - | - | - | - |
| | a | 6.2 | 26 | 48.8 | 2 | 12 | 2 | 14 | 7 | 1 | 11 | 6 | 4 | * | * | - | 12 | 2 | 1 | 1 | - | 4 | - | 10 |
| (N2)PKNaMg | b | 5.7 | 26 | 45.8 | 6 | 18 | 10 | 11 | 4 | 4 | 15 | 5 | 3 | - | 2 | * | 10 | * | 1 | * | - | 5 | - | 2 |
| (9/1) | c | 4.6 | 24 | 28.4 | 42 | 1 | 18 | 1 | * | 19 | 3 | 2 | 2 | * | - | - | 1 | * | 1 | * | - | 5 | - | 5 |
| | d | 3.9 | 7 | 31.9 | 13 | - | 72 | - | - | * | 15 | - | - | - | - | - | - | - | - | - | - | * | - | - |
| | a | 6.9 | 26 | 70.6 | * | 18 | - | 45 | 5 | 3 | * | 2 | 2 | * | - | - | 2 | * | 3 | 1 | - | * | - | * |
| N*2PKNaMg | b | 6.6 | 23 | 78.0 | * | 21 | * | 39 | 5 | 4 | * | 2 | 5 | * | - | - | 2 | - | 3 | 1 | - | * | - | * |
| (14/2) | c | 6.0 | 23 | 72.7 | * | 19 | * | 36 | 5 | 3 | 1 | 3 | 7 | * | - | - | 4 | - | 5 | 2 | - | * | - | - |
| | d | 5.9 | 23 | 79.4 | * | 23 | * | 26 | 9 | 1 | 3 | 2 | 8 | * | - | - | 2 | * | 4 | 2 | - | * | - | - |
| | a | 6.8 | 29 | 55.9 | 1 | 6 | * | 17 | 3 | 9 | * | 3 | 3 | * | * | - | 3 | 10 | 1 | 6 | - | 5 | * | 19 |
| (N*2)PKNaMg | b | 6.3 | 28 | 52.5 | 1 | 8 | 1 | 15 | 3 | 11 | 8 | 5 | 7 | * | * | - | 4 | 7 | 1 | 6 | * | 2 | - | 14 |
| (14/1) | c | 5.7 | 28 | 47.4 | 2 | 6 | 2 | 12 | 4 | 3 | 11 | 4 | 10 | - | 1 | - | 4 | 8 | 1 | 9 | - | 6 | - | 12 |
| | d | 5.7 | 24 | 54.1 | 4 | 5 | 5 | 9 | 4 | 2 | 10 | 4 | 12 | - | - | * | 3 | 13 | 1 | 9 | - | 7 | - | 11 |
| | a | 6.1 | 17 | 80.5 | * | 20 | - | 37 | 10 | * | 9 | 1 | 5 | - | - | - | * | - | 3 | * | - | - | - | - |
| N3PKNaMg | b | 5.8 | 16 | 72.5 | * | 22 | * | 32 | 18 | * | 13 | 2 | 4 | - | - | - | 1 | - | 1 | - | - | - | - | - |
| (11/1) | c | 4.6 | 15 | 64.1 | 7 | 33 | 2 | 15 | 5 | * | 33 | 1 | 1 | - | - | - | * | - | 1 | - | - | - | - | - |
| | d | 3.5 | 2 | 53.3 | - | - | * | - | - | - | 100 | - | - | - | - | - | - | - | - | - | - | - | - | - |
| | a | 6.4 | 16 | 74.9 | * | 20 | * | 38 | 10 | * | 4 | * | 7 | - | - | - | * | - | 2 | - | - | * | - | - |
| N3PKNaMgSi | b | 5.6 | 12 | 79.2 | 5 | 37 | - | 31 | 10 | - | 13 | * | 1 | - | - | - | * | - | * | - | - | - | - | - |
| (11/2) | c | 4.7 | 14 | 57.9 | 10 | 45 | 1 | 27 | 1 | * | 12 | 2 | 1 | - | - | - | * | - | * | - | - | - | - | - |
| | d | 3.6 | 3 | 69.0 | 3 | * | - | - | - | - | 97 | - | - | - | - | - | - | - | - | - | - | - | - | - |

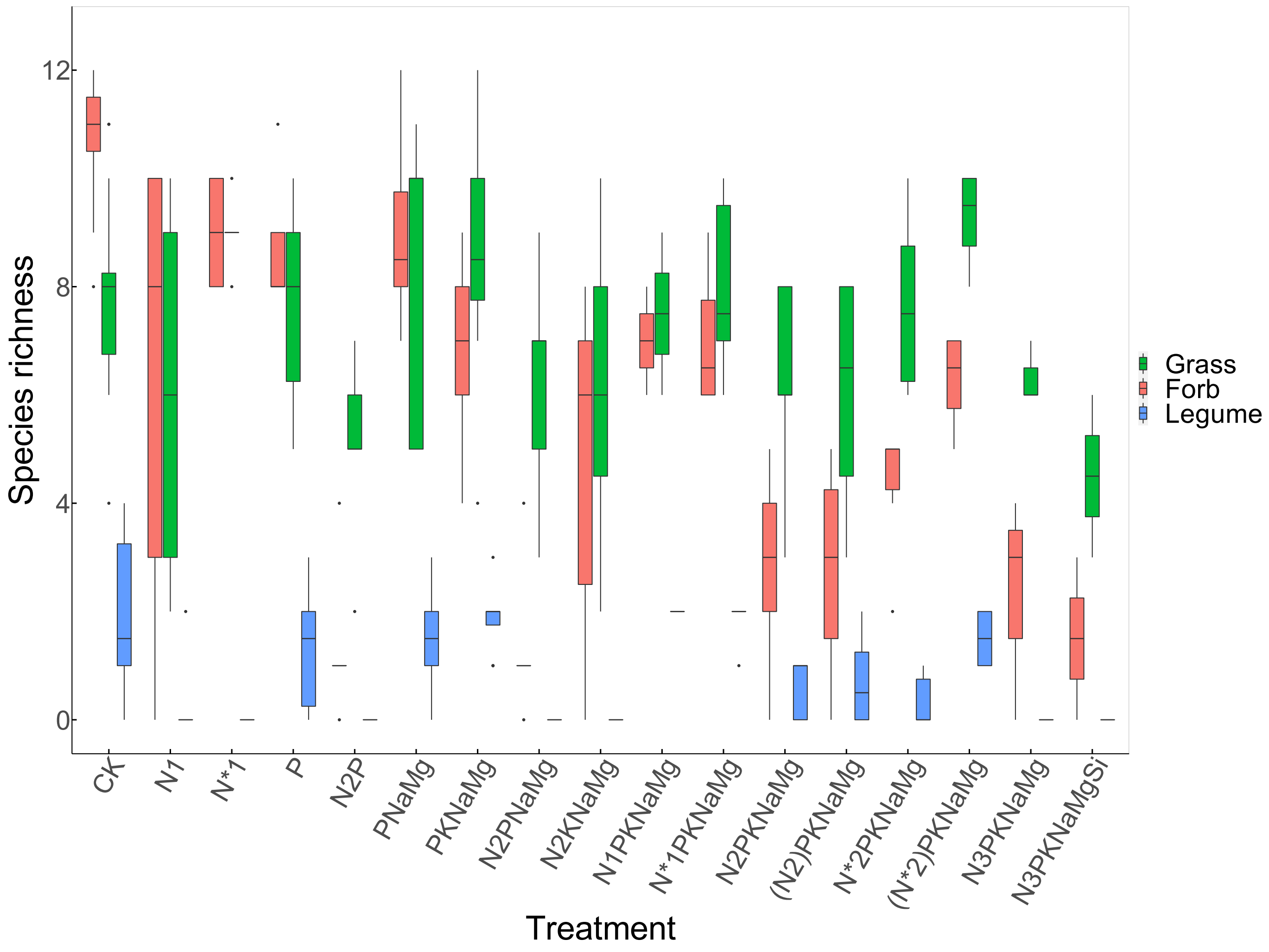


**Fig. S1** Species richness of different functional groups (FG: grass, forb and legume) across the 15 different treatments of nutrient addition crossed with different levels of soil pH adjustments. Please note (N2)PKNaMg and (N*)2PKNaMg indicate N addition is withheld in one half of N2PKNaMg and N*2PKNaMg respectively since 1990.

**Appendix S2 Details of model fitting and cross-validation**

*Model fitting*

Since the experimental treatments in the Park Grass have "diversified" several times over 150 years (Appendix S1), to make the best use of the rich data, we organized all the time-series data into two subsets and fitted Lotka-Volterra competition models separately for each subset. The first "recent" subset is the time-series data of all the plots from 1965 to 2000 in which four levels of soil pH treatment (unlimed, 5, 6 and 7) were applied in each treatment of nutrient addition. The second "longul" subset is the time-series data of unlimed ("U") vs limed ("L") plots since 1856, which include all plots from 1862 to 1902 (regarded as "unlimed", given the data points all lie before the small amount of lime was applied in the late 1880s), all unlimed and limed half-plots from 1903 to 1964, and all unlimed ("d") and limed ("a", "b" or "c", depending on the similarity of its pH to that of the corresponding limed half-plots before 1965) quarter-plots from 1965 to 2000. For more details of these plots with various treatments, please see Appendix S1. Note except for all plots from 1991 to 2000 with six replicates per plot, all the other plots have one replicate per plot. Moreover, it becomes technically impossible to fit models with a species when it occurs in less than 30% of all data points or contributes little biomass in a plot, so we merge these species as "rare species" to facilitate the fitting. In this way, the number of species *explicitly* fitted in the models is 13 on average, but these species comprise > 95% of total biomass in a plot (Table S3). In other words, although there are a number of "rare species" in some plots, they only comprise a tiny proportion of total biomass (Table S3). So, even if we completely removed the "rare species" from fitted models, the results remained very similar for the explicitly fitted species (not shown). Moreover, we also tried to use the criteria of 40% and 50% to fit models but the estimates of the parameters ($r$ and $\alpha$, i.e. intrinsic population growth rates and competition coefficients) were very similar to those of 30% (Fig. S2). Therefore, we decided to apply the 30% criteria in order to include as many species as possible but still keep the adequate reliability of model fittings.

Separately for each subset, we fitted ln-transformed Lotka-Volterra competition models with time-series data of each plot using Bayesian inference with the “rstan” package. In the fittings, we used uninformative priors for unknown parameters $r_i$, $\alpha_{ii}$ and $\alpha_{ij}$, and used normally distributed residuals for all models (y ~ normal (ŷ, sigmaeps)). We assumed positive $r$ for all the modelled species, for two reasons. First, most of the species we *explicitly* modelled in both the “recent” and “longul” subsets are necessarily extant throughout the respective time-series, so their $r$ are reasonably positive (otherwise, they would have disappeared). Second, in the “longul” subset, there are indeed a small number of species that pass the 30% criteria but their $r$ might be negative since they occurred in earlier times but have disappeared over time. These species mainly occur in the unlimed (“U”) plots of treatments with the addition of N as $NH_4$, and importantly the model fitting of these plots was generally poor, so we had to drop them from the further analyses (Table S3). Moreover, we assumed all the $\alpha_{ii}$ and $\alpha_{ij}$ are positive, i.e. net interactions between species are assumed overall competitive, although facilitative effects might also partially occur between some species (e.g. legumes and other species). For each model, we ran four chains, each with 3500 iterations in which 1000 iterations were discarded as “warmup” while the remaining 2500 iterations were drawn as posterior samples. The detailed diagnostics of model fits are provided in Table S3. The $n_{eff}$ is the mean size of effective samples of the parameters ($r$ and $\alpha$), and the $\hat{R}$ is the mean potential scale reduction factor of the parameters that indicates the degree of convergence (greater deviation from 1 implies more severe convergence problems). From the values of both $n_{eff}$ and $\hat{R}$ and their SDs, it is reasonable to conclude that the model fits are overall adequately reliable. Finally, the correlations between intrinsic population growth rates ($r_i$) and intraspecific and interspecific competition coefficients ($a_{ii}$, $a_{ij}$) across all species and treatments are negligible (Fig. S3), which indicates the independence in estimation of these parameters.

*Cross-validation*

Following the suggestions of Tredennick et al. (2017), we applied the "leave-one-year-out" cross-validation to further assess the goodness-of-fit of all the models. Specifically, for all species in a plot, we first fitted models with time-series data from all years except one, and then used the fitted models to predict biomasses in the removed year from observed biomasses in the previous year ("out-of-sample" prediction). In order to include the random variable (i.e. "period", see the main text) in prediction, we used the random effect of the corresponding period estimated from the model fitted with data from all years (because the random effect of the very period simply cannot be estimated if the year is removed). For example, if 1994 is left out, it becomes impossible to estimate the random effect of the 1994-1995 period that is nevertheless required for the subsequent prediction, so we use the random effect of the 1994-1995 period estimated from the model fitted with data from all years. We repeated the cross-validation procedure for all possible years. Please note this means we could only do the cross-validation for the years from 1991 to 2000 for both the "recent" and "longul" subsets (although the models were fitted with the whole time-series), simply because the "leave-one-year-out" is impossible for the years before 1991 where the data are often collected every ten to twenty years. After predictions, we calculated Pearson correlations ($\rho$) between predictions and observations and absolute error (i.e. |predicted - observed|) to measure the prediction accuracy and error of fitted models across species, plots and years (from 1991 to 2000) (Fig. S4). Based on this, we removed a couple of species (e.g. *Helictotrichon pubescens*) from a few plots (e.g. the 14 "U" plot) because of their poor accuracies or high errors. The relatively high median accuracy (recent: 0.50; longul: 0.60) and low median absolute error (recent: 1.7; longul: 0.88; unit: g/0.125 $m^2$) (Fig. S4) indicate the reliable goodness-of-fit of the fitted models. After all this, we obtained a total of 36 explicitly modelled species and 11455 species pairs (CK: 2883; fertilised: 8572) across all plots and treatments for further analyses.

**Table S3** Species richness and relative abundance of explicitly fitted species ($SR_{fit}$ and $RA_{fit}$) and "rare species" ($SR_{rare}$ and $RA_{rare}$), and posterior summary statistics of the parameters from Lotka-Volterra competition models fitted with time-series data of the "recent" (1965-2000) and "longul" (1856-2000) subsets, in each of the sub-plots (SP: a, b, c and d for the "recent" subset, and U and L for the "longul" subset) within each of the plots with 15 different treatments of nutrient addition. Note all of U subplots with $NH_4^+$ (i.e. $(NH_4)_2SO_4$) are dropped because of the poor model fitting and cross-validation. The mean number of effective samples ($n_{eff}$) and mean potential scale reduction factor ($\hat{R}$) (deviation from 1 indicates convergence problems) and their SDs are calculated across the parameters ($r$ and $\alpha$).

| Treatment (plot) | SP | $SR_{fit}$ | $SR_{rare}$ | $RA_{fit}$ | $RA_{rare}$ | $n_{eff}$ | SD ($n_{eff}$) | $\hat{R}$ | SD ($\hat{R}$) |
|---|---|---|---|---|---|---|---|---|---|
| CK (i.e. Nil) (2/2, 3, 12) | a | 24 | 21 | 95.7 | 4.3 | 3703 | 529 | 1.001 | 0.0007 |
| | b | 22 | 21 | 94.3 | 5.7 | 4105 | 628 | 1.001 | 0.0005 |
| | c | 17 | 21 | 96.0 | 4.0 | 4969 | 939 | 1.000 | 0.0005 |
| | d | 14 | 25 | 94.7 | 5.3 | 4307 | 954 | 1.001 | 0.0006 |
| | U | 22 | 40 | 94.7 | 5.3 | 11551 | 2012 | 0.999 | 0.0002 |
| | L | 25 | 38 | 95.0 | 5.0 | 11181 | 2054 | 0.999 | 0.0002 |
| N1 (1) | a | 19 | 16 | 95.2 | 4.8 | 4622 | 570 | 1.001 | 0.0005 |
| | b | 14 | 16 | 92.2 | 7.8 | 4602 | 405 | 1.000 | 0.0004 |
| | c | 6 | 18 | 96.0 | 4.0 | 4120 | 506 | 1.000 | 0.0004 |
| | d | 2 | 3 | 98.4 | 1.6 | 1780 | 1008 | 1.002 | 0.0027 |
| | L | 22 | 26 | 93.3 | 6.7 | 11719 | 1162 | 0.999 | 0.0001 |
| N*1 (17) | a | 17 | 19 | 94.0 | 6.0 | 4442 | 1020 | 1.001 | 0.0008 |
| | b | 17 | 21 | 95.5 | 4.5 | 5067 | 1177 | 1.000 | 0.0004 |
| | c | 16 | 18 | 96.8 | 3.2 | 4394 | 789 | 1.000 | 0.0004 |
| | d | 17 | 19 | 95.0 | 5.0 | 4119 | 833 | 1.001 | 0.0007 |
| | U | 20 | 41 | 93.4 | 6.6 | 13987 | 1230 | 1.000 | 0.0001 |
| | L | 18 | 42 | 93.6 | 6.4 | 13483 | 1436 | 1.000 | 0.0002 |
| P (4/1) | a | 20 | 17 | 95.8 | 4.2 | 3992 | 765 | 1.001 | 0.0003 |
| | b | 22 | 13 | 95.4 | 4.6 | 3542 | 656 | 1.001 | 0.0007 |
| | c | 14 | 19 | 92.8 | 7.2 | 4803 | 958 | 1.000 | 0.0004 |
| | d | 13 | 24 | 94.3 | 5.7 | 4358 | 703 | 1.000 | 0.0006 |
| | U | 17 | 41 | 95.0 | 5.0 | 10726 | 1647 | 1.000 | 0.0003 |
| | L | 27 | 29 | 98.3 | 1.7 | 10041 | 1939 | 1.000 | 0.0002 |
| N2P (4/2) | a | 10 | 14 | 88.1 | 11.9 | 3764 | 747 | 1.001 | 0.0008 |
| | b | 6 | 10 | 91.0 | 9.0 | 3912 | 654 | 1.000 | 0.0004 |

| | | | | | | | | | |
|---|---|---|---|---|---|---|---|---|---|
| | c | 6 | 12 | 99.3 | 0.7 | 4148 | 638 | 1.000 | 0.0005 |
| | d | 2 | 6 | 98.9 | 1.1 | 3107 | 896 | 1.001 | 0.0009 |
| | L | 8 | 32 | 98.6 | 1.4 | 10698 | 1604 | 1.000 | 0.0002 |
| PNaMg (8) | a | 21 | 19 | 94.5 | 5.5 | 4637 | 1036 | 1.001 | 0.0008 |
| | b | 21 | 17 | 95.8 | 4.2 | 2818 | 331 | 1.001 | 0.0007 |
| | c | 13 | 20 | 92.7 | 7.3 | 5047 | 765 | 1.001 | 0.0005 |
| | d | 13 | 20 | 95.5 | 4.5 | 4132 | 788 | 1.001 | 0.0006 |
| | U | 23 | 35 | 97.6 | 2.4 | 13775 | 1310 | 1.000 | 0.0001 |
| | L | 28 | 29 | 98.4 | 1.6 | 12355 | 1477 | 1.000 | 0.0002 |
| PKNaMg (7, 15) | a | 19 | 10 | 97.3 | 2.7 | 3652 | 641 | 1.001 | 0.0007 |
| | b | 19 | 12 | 96.3 | 3.7 | 4138 | 793 | 1.001 | 0.0008 |
| | c | 15 | 16 | 94.2 | 5.8 | 3774 | 775 | 1.001 | 0.0005 |
| | d | 13 | 15 | 94.8 | 5.2 | 3681 | 983 | 1.001 | 0.0011 |
| | U | 20 | 36 | 97.0 | 3.0 | 12052 | 1564 | 1.000 | 0.0002 |
| | L | 21 | 32 | 90.5 | 9.5 | 11149 | 1649 | 1.000 | 0.0002 |
| N2PNaMg (10) | a | 11 | 12 | 98.1 | 1.9 | 3473 | 533 | 1.001 | 0.0006 |
| | b | 8 | 7 | 99.5 | 0.5 | 3289 | 858 | 1.001 | 0.0007 |
| | c | 6 | 8 | 97.1 | 2.9 | 2630 | 612 | 1.001 | 0.0009 |
| | d | 3 | 0 | 100.0 | 0.0 | 2306 | 1740 | 1.003 | 0.0022 |
| | L | 10 | 28 | 98.4 | 1.6 | 13118 | 575 | 1.000 | 0.0002 |
| N2KNaMg (18) | a | 12 | 21 | 89.2 | 10.8 | 4489 | 471 | 1.000 | 0.0004 |
| | b | 9 | 23 | 89.5 | 10.5 | 4177 | 495 | 1.000 | 0.0004 |
| | c | 7 | 16 | 82.2 | 17.8 | 3640 | 400 | 1.001 | 0.0005 |
| | d | 2 | 4 | 97.2 | 2.8 | 2255 | 362 | 1.001 | 0.0008 |
| | L | 17 | 27 | 93.1 | 6.9 | 7965 | 1314 | 1.000 | 0.0002 |
| N1PKNaMg (6) | a | 19 | 9 | 98.9 | 1.1 | 5206 | 796 | 1.000 | 0.0003 |
| | b | 15 | 12 | 91.5 | 8.5 | 4967 | 609 | 1.000 | 0.0006 |
| N*1PKNaMg (16) | a | 18 | 8 | 95.0 | 5.0 | 5442 | 701 | 1.000 | 0.0004 |
| | b | 15 | 11 | 92.4 | 7.6 | 4226 | 723 | 1.000 | 0.0003 |
| | c | 16 | 12 | 95.4 | 4.6 | 3653 | 867 | 1.001 | 0.0008 |
| | d | 16 | 10 | 97.4 | 2.6 | 3299 | 913 | 1.001 | 0.0007 |
| | U | 19 | 28 | 95.2 | 4.8 | 11032 | 2112 | 1.000 | 0.0002 |
| | L | 21 | 26 | 96.7 | 3.3 | 11417 | 1443 | 1.000 | 0.0001 |
| N2PKNaMg (9/2) | a | 13 | 11 | 96.5 | 3.5 | 4101 | 621 | 1.001 | 0.0007 |
| | b | 11 | 8 | 93.7 | 6.3 | 4382 | 1493 | 1.001 | 0.0015 |
| | c | 9 | 9 | 85.7 | 14.3 | 3449 | 720 | 1.001 | 0.0006 |
| | d | 3 | 1 | 100.0 | 0.0 | 2650 | 1019 | 1.001 | 0.0003 |
| | L | 12 | 32 | 94.8 | 5.2 | 9010 | 1789 | 1.000 | 0.0002 |
| (N2)PKNaMg (9/1) | a | 16 | 10 | 96.8 | 3.2 | 3596 | 820 | 1.001 | 0.0007 |
| | b | 13 | 13 | 94.2 | 5.8 | 3382 | 951 | 1.001 | 0.0009 |
| | c | 7 | 17 | 86.4 | 13.6 | 2757 | 898 | 1.002 | 0.0020 |
| | d | 3 | 4 | 99.5 | 0.5 | 2543 | 302 | 1.001 | 0.0006 |
| N*2PKNaMg | a | 10 | 16 | 95.9 | 4.1 | 3847 | 873 | 1.001 | 0.0005 |

| | | | | | | | | | |
|---|---|---|---|---|---|---|---|---|---|
| (14/2) | b | 11 | 12 | 97.7 | 2.3 | 4494 | 636 | 1.001 | 0.0004 |
| | c | 12 | 11 | 97.6 | 2.4 | 3602 | 599 | 1.001 | 0.0004 |
| | d | 13 | 10 | 98.4 | 1.6 | 4585 | 638 | 1.000 | 0.0004 |
| | U | 14 | 26 | 90.4 | 9.6 | 11038 | 2619 | 1.000 | 0.0002 |
| | L | 16 | 25 | 98.0 | 2.0 | 11300 | 1429 | 1.000 | 0.0003 |
| (N*2)PKNaMg (14/1) | a | 18 | 11 | 96.7 | 3.3 | 3822 | 823 | 1.000 | 0.0006 |
| | b | 17 | 11 | 96.6 | 3.4 | 4487 | 689 | 1.000 | 0.0005 |
| | c | 18 | 10 | 97.2 | 2.8 | 5281 | 932 | 1.000 | 0.0005 |
| | d | 17 | 7 | 98.9 | 1.1 | 4192 | 927 | 1.001 | 0.0005 |
| N3PKNaMg (11/1) | a | 10 | 7 | 99.5 | 0.5 | 3879 | 563 | 1.001 | 0.0006 |
| | b | 9 | 7 | 98.6 | 1.4 | 3690 | 598 | 1.001 | 0.0006 |
| | c | 7 | 8 | 97.0 | 3.0 | 3258 | 1062 | 1.002 | 0.0028 |
| | d | 1 | 1 | 99.9 | 0.1 | 3176 | 229 | 1.002 | 0.0002 |
| | L | 11 | 5 | 99.4 | 0.6 | 9521 | 2210 | 1.000 | 0.0004 |
| N3PKNaMgSi (11/2) | a | 7 | 9 | 97.0 | 3.0 | 4067 | 549 | 1.001 | 0.0006 |
| | b | 5 | 7 | 91.0 | 9.0 | 2842 | 795 | 1.001 | 0.0008 |
| | c | 5 | 9 | 96.3 | 3.7 | 2313 | 1308 | 1.019 | 0.0409 |
| | d | 1 | 2 | 96.9 | 3.1 | 3116 | 275 | 1.001 | 0.0002 |
| | L | 10 | 8 | 99.5 | 0.5 | 9462 | 2332 | 1.000 | 0.0002 |

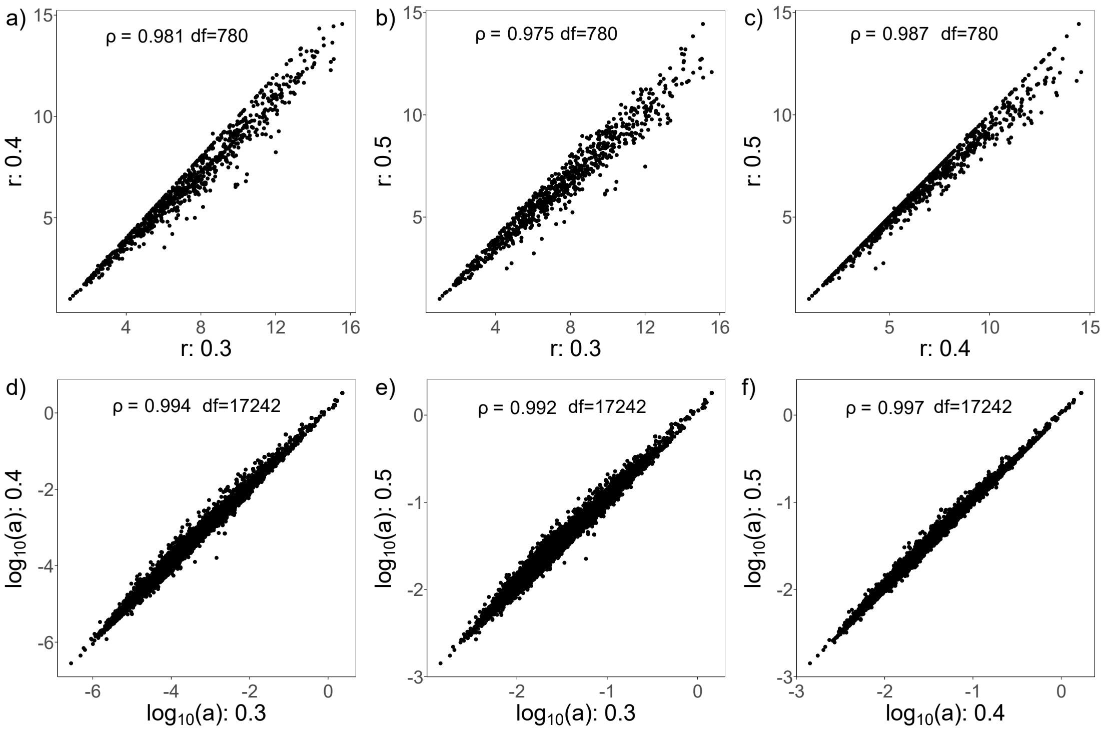


**Fig. S2** Pearson correlations ($\rho$) between $r$ (intrinsic population growth rate: a, b and c) or $\alpha$ (intraspecific and interspecific competition coefficients: d, e and f) derived from Lotka-Volterra competition models fitted with time-series data based on the selection criteria of 30%, 40% and 50% (see the text above).

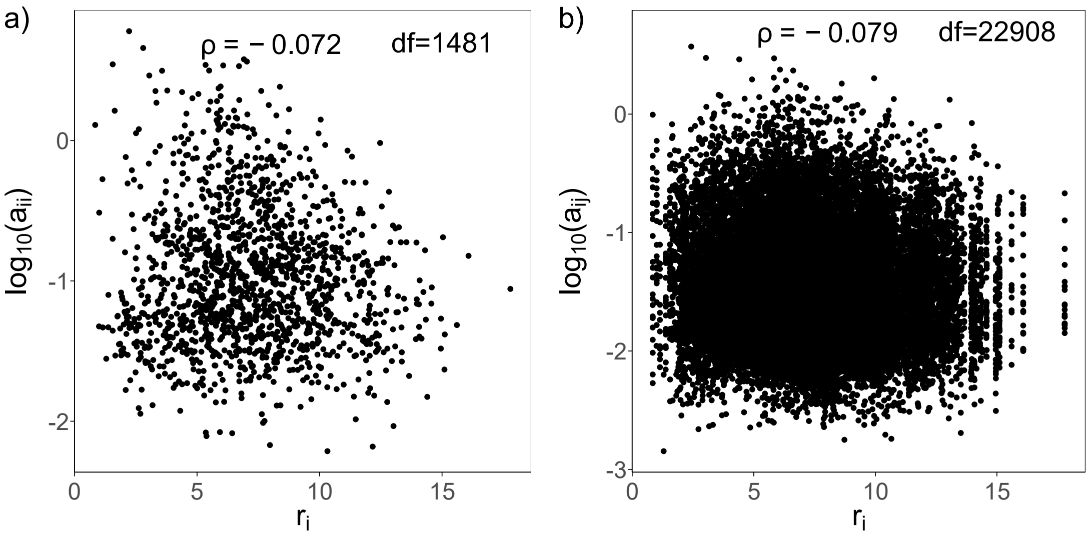


**Fig. S3** Pearson correlations ($\rho$) between intrinsic population growth rates ($r_i$) and a) intraspecific competition coefficients ($a_{ii}$), and b) interspecific competition coefficients ($a_{ij}$) across all species and treatments.

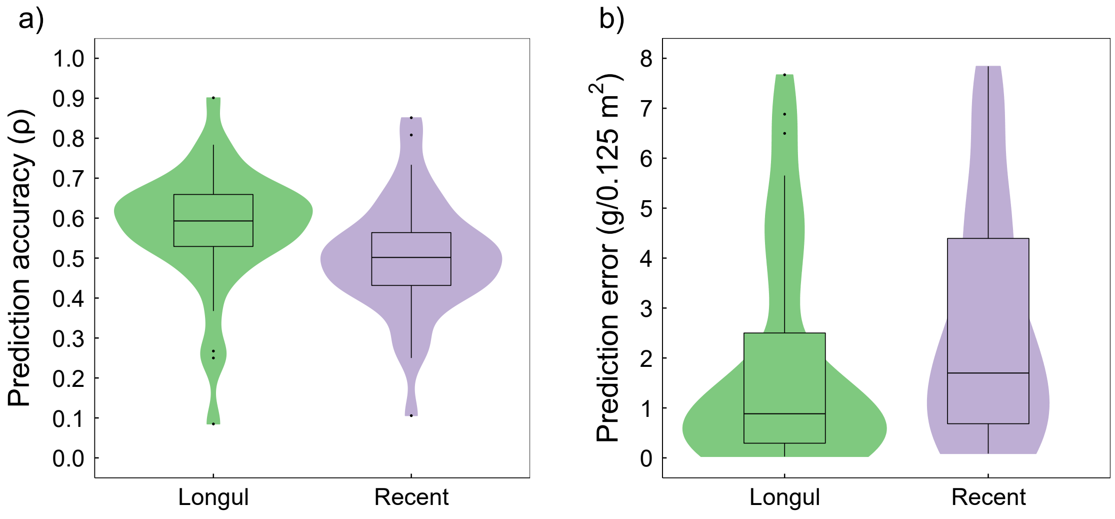


**Fig. S4** Prediction accuracies (i.e. Pearson correlation between predictions and observations, $\rho$) (a) and absolute error (i.e. |predicted - observed|) (b) of fitted models in the "leave-one-year-out" cross-validation for the "recent" subset (1965-2000) and the "longul" subset (1856-2000).

### Appendix S3 Supplementary results of the main analyses

This appendix includes the additional important results of the main analyses that could not be elaborated on in the main text. Briefly, Table S4 shows the detailed statistical results of linear models testing the effects of different treatments of nutrient addition and soil pH, “functional group” and their interactions on intrinsic population growth rate ($r$), niche difference (ND: $1-\sqrt{\frac{\alpha_{ij}\alpha_{ji}}{\alpha_{ii}\alpha_{jj}}}$), competitive difference (CD: $\sqrt{\frac{\alpha_{jj}\alpha_{ji}}{\alpha_{ii}\alpha_{ij}}}$), and per-capita intraspecific and interspecific competition ($a_{ii}$ and $a_{ij}$), and Fig. S5 illustrates all the significant and marginally significant effects in the models. Table S5 shows the detailed statistical results of linear models testing the effects of N withholding and other important covariates and their interactions on $r$, ND, CD, $a_{ii}$ and $a_{ij}$. Please remember that withholding N addition was introduced in half of the N2PKNaMg ($NH_4^+$) and N*2PKNaMg ($NO_3^-$) treatments since 1990 to test whether the withholding could make the community diversity and composition (which have changed greatly due to the treatments) recover. Table S6 shows the competitive hierarchy of all explicitly fitted species (constructed based on competitive differences between species) in each of the plots with various treatments of nutrient addition and soil pH. Table S7 shows the detailed statistical results of generalized linear models testing the effects of the changes (due to the various treatments of nutrient addition and soil pH) in plot-level $r$, ND, absolute CD (absolute values of CD, reflecting absolute competitive asymmetries between species) and their interactions on the plot-level species richness. For the details of constructing the abovementioned models, see the *Statistical analyses* in the main text.

**Table S4** Results of linear models testing the effects of N type (0, $NH_4^+$, $NO_3^-$), P (0, 1), K (0, 1), NaMg (0, 1), soil pH, "functional group (FG)" and their interactions on intrinsic population growth rate ($r$), niche difference (ND: $1-\sqrt{\frac{\alpha_{ij}\alpha_{ji}}{\alpha_{ii}\alpha_{jj}}}$), competitive difference (CD: $\sqrt{\frac{\alpha_{jj}\alpha_{ji}}{\alpha_{ii}\alpha_{ij}}}$), and per-capita intraspecific and interspecific competition ($a_{ii}$ and $a_{ij}$). Note that all the variables, except for $r$ (square root transformed) and ND, are log10-transformed to meet the assumptions for residuals. Up and down arrows beside significant (p<0.05, in bold) or marginally significant (0.05⩽p<0.1, in italic) p values indicate positive and negative effects respectively.

| **Variables** | | ***r*** | | **ND** | | **CD** | | $\boldsymbol{a_{ii}}$ | | $\boldsymbol{a_{ij}}$ | |
|---|---|---|---|---|---|---|---|---|---|---|---|
| | df | χ² | p | χ² | p | χ² | p | χ² | p | χ² | p |
| N type | 2 | 111.55 | **<0.001**↓ | 14.87 | **0.001**↑ | 14.33 | **0.001**↓ | 7.08 | **0.029**↓ | 64.53 | **<0.001**↓ |
| P | 1 | 11.84 | **0.001**↑ | 4.28 | **0.039**↑ | 65.13 | **<0.001**↓ | 23.06 | **<0.001**↓ | 340.80 | **<0.001**↓ |
| K | 1 | 7.78 | **0.005**↑ | 7.79 | **0.005**↓ | 1.21 | 0.271 | 9.75 | **0.002**↓ | 192.76 | **<0.001**↓ |
| NaMg | 1 | 4.68 | **0.031**↓ | 3.08 | *0.079*↓ | 4.77 | **0.029**↑ | 0.64 | 0.424 | 1.38 | 0.241 |
| pH | 1 | 287.20 | **<0.001**↑ | 17.38 | **<0.001**↓ | 25.85 | **<0.001**↓ | 20.28 | **<0.001**↓ | 354.67 | **<0.001**↓ |
| FG | 2/5/8 | 19.98 | **<0.001** | 59.82 | **<0.001** | 109.19 | **<0.001** | 107.97 | **<0.001** | 1372.63 | **<0.001** |
| N type: P | 1 | 11.73 | **0.001** | 5.92 | **0.015** | 0.35 | 0.556 | 0.76 | 0.383 | 1.100 | 0.294 |
| N type: K | 1 | 0.72 | 0.398 | 1.61 | 0.205 | 1.81 | 0.179 | 0.61 | 0.436 | 10.568 | **0.001** |
| N type: NaMg | 1 | 2.64 | 0.104 | 2.24 | 0.135 | 0.90 | 0.342 | 2.91 | *0.088* | 19.445 | **<0.001** |
| P: K | 1 | 24.63 | **<0.001** | 30.15 | **<0.001** | 0.08 | 0.781 | 3.89 | **0.049** | 17.729 | **<0.001** |
| pH: N type | 2 | 116.95 | **<0.001** | 6.15 | **0.046** | 20.56 | **<0.001** | 6.27 | **0.044** | 69.301 | **<0.001** |
| pH: P | 1 | 7.42 | **0.006** | 3.75 | *0.053* | 9.51 | **0.002** | 0.93 | 0.334 | 33.003 | **<0.001** |
| pH: K | 1 | 0.29 | 0.588 | 1.16 | 0.282 | 6.72 | **0.010** | 1.26 | 0.262 | 20.077 | **<0.001** |
| pH: NaMg | 1 | 6.27 | **0.012** | 1.23 | 0.268 | 0.74 | 0.388 | 0.07 | 0.793 | 0.442 | 0.506 |
| FG: N type | 4/10/16 | 7.54 | 0.110 | 27.19 | **0.002** | 39.79 | **0.001** | 11.80 | **0.019** | 165.227 | **<0.001** |
| FG: P | 2/5/8 | 10.68 | **0.005** | 55.03 | **<0.001** | 45.84 | **<0.001** | 1.69 | 0.430 | 10.704 | 0.219 |
| FG: K | 2/5/8 | 0.65 | 0.722 | 19.74 | **0.001** | 24.35 | **0.002** | 6.46 | **0.040** | 113.472 | **<0.001** |
| FG: NaMg | 2/5/8 | 0.99 | 0.610 | 41.59 | **<0.001** | 43.35 | **<0.001** | 3.15 | 0.207 | 81.884 | **<0.001** |
| FG: pH | 2/5/8 | 15.46 | **<0.001** | 22.19 | **<0.001** | 96.71 | **<0.001** | 20.03 | **<0.001** | 182.935 | **<0.001** |

**Fig. S5** The plots illustrating all the significant and marginally significant main and interaction effects in the linear models (see Table S4) testing the effects of N type (0, $NH_4^+$, $NO_3^-$), P (0, 1), K (0, 1), NaMg (0, 1), soil pH, "functional group (FG)" and their interactions on intrinsic population growth rate ($r$; Fig. S5.1), niche difference (ND: $1-\sqrt{\frac{\alpha_{ij}\alpha_{ji}}{\alpha_{ii}\alpha_{jj}}}$; Fig. S5.2), competitive difference (CD: $\sqrt{\frac{\alpha_{jj}\alpha_{ji}}{\alpha_{ii}\alpha_{ij}}}$; Fig. S5.3), and per-capita intraspecific and interspecific competition ($a_{ii}$ and $a_{ij}$; Fig. S5.4 and S5.5). Note that all the variables, except for $r$ (square root transformed) and ND, are $\log_{10}$-transformed to meet the assumptions for residuals.

**Fig. S5.1** Intrinsic popolation growth rate (r)

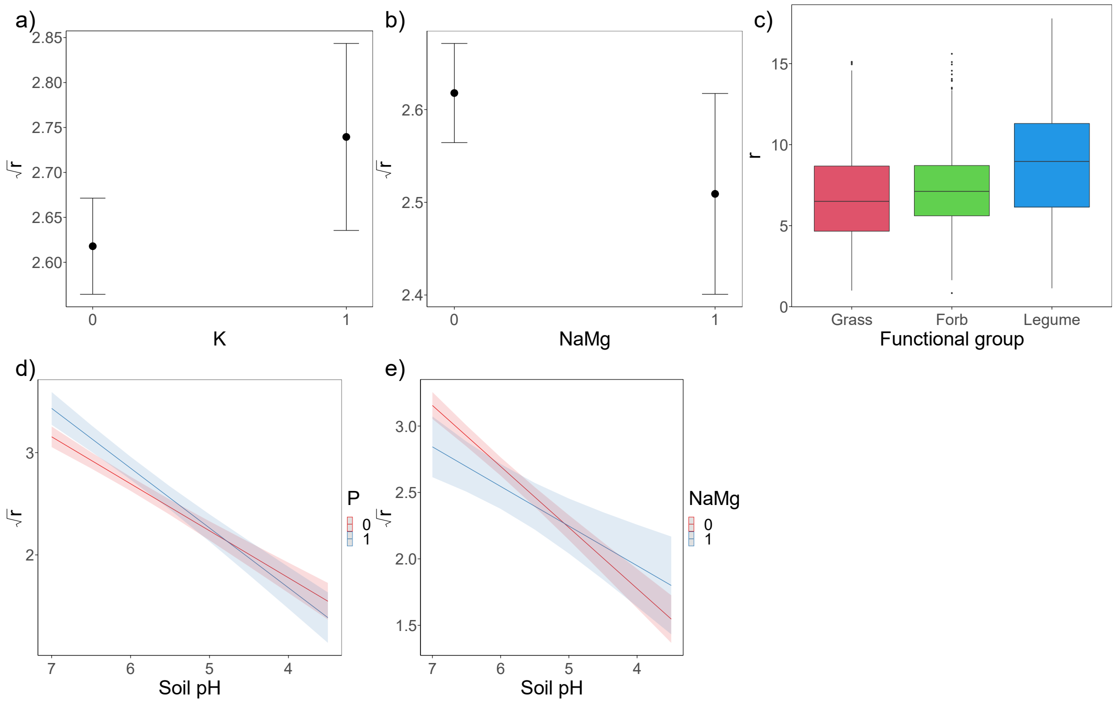

**Fig. S5.2**

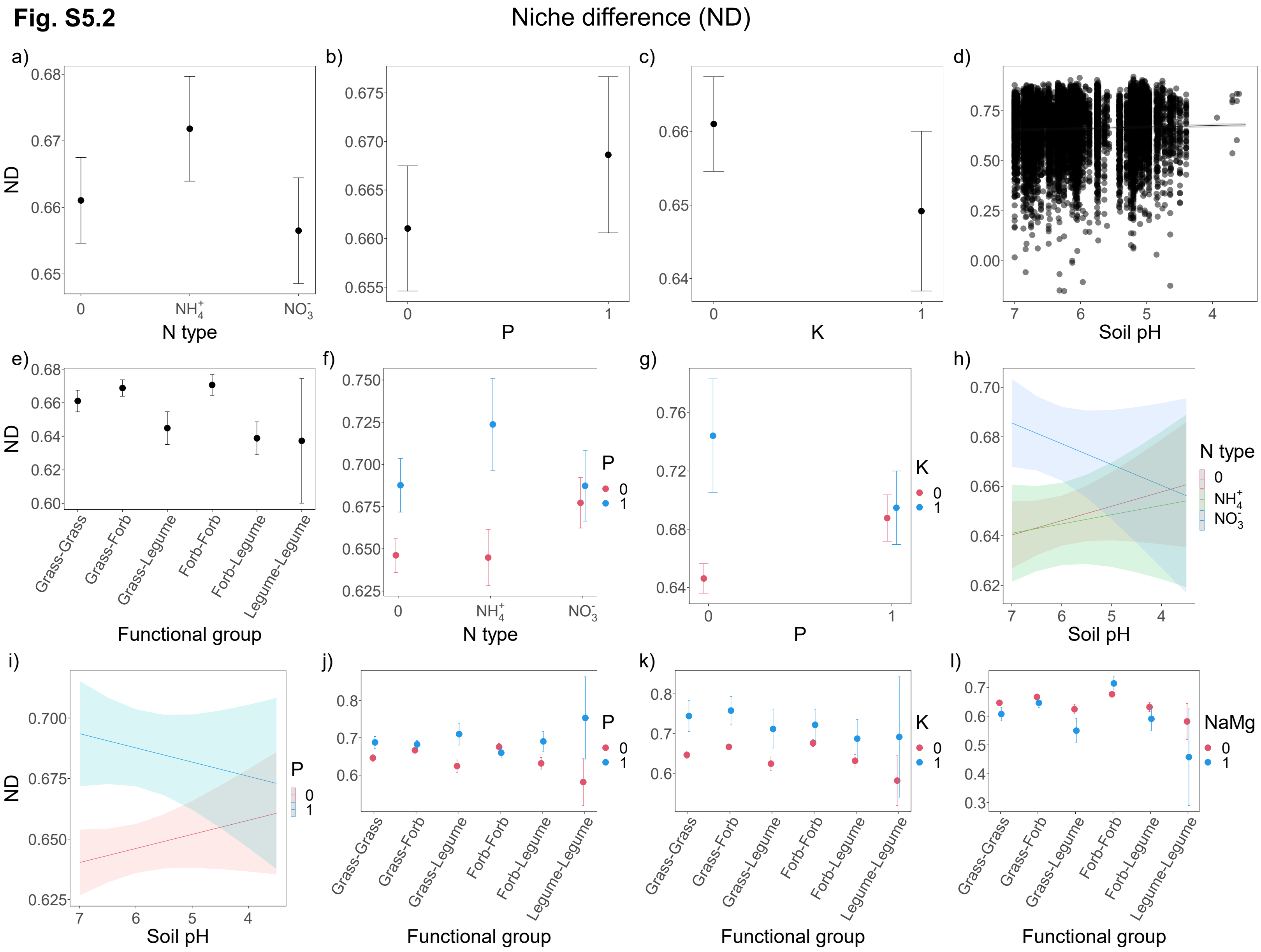

Niche difference (ND)
a)
b)
c)
d)
e)
f)
g)
h)
i)
j)
k)
l)
ND
N type
P
K
Soil pH
Functional group
Grass-Grass
Grass-Forb
Grass-Legume
Forb-Forb
Forb-Legume
Legume-Legume
$NH_4^+$
$NO_3^-$
NaMg

**Fig. S5.3**

Competitive difference (CD)

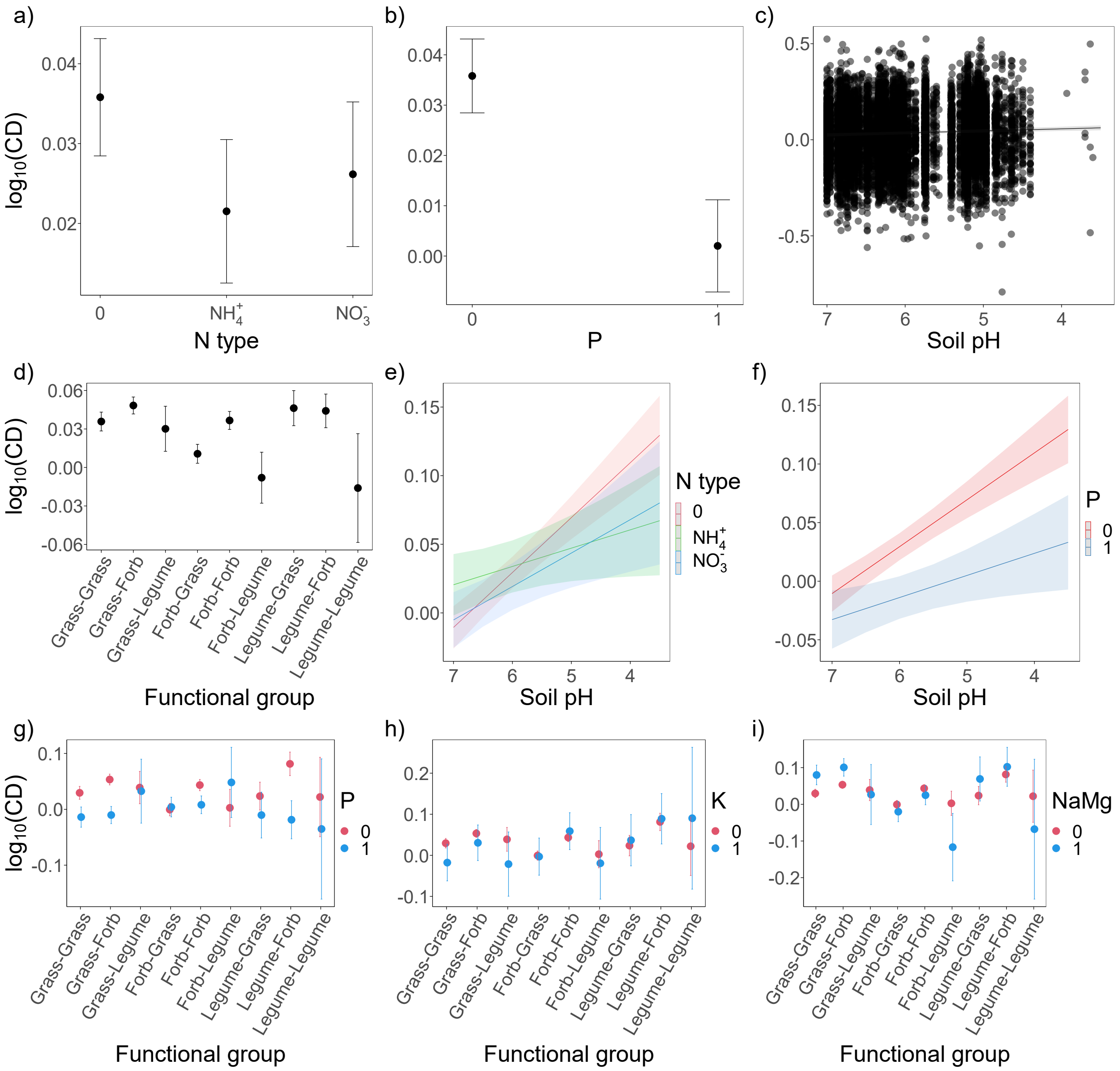

**Fig. S5.4**

Intraspecific competition ($a_{ii}$)

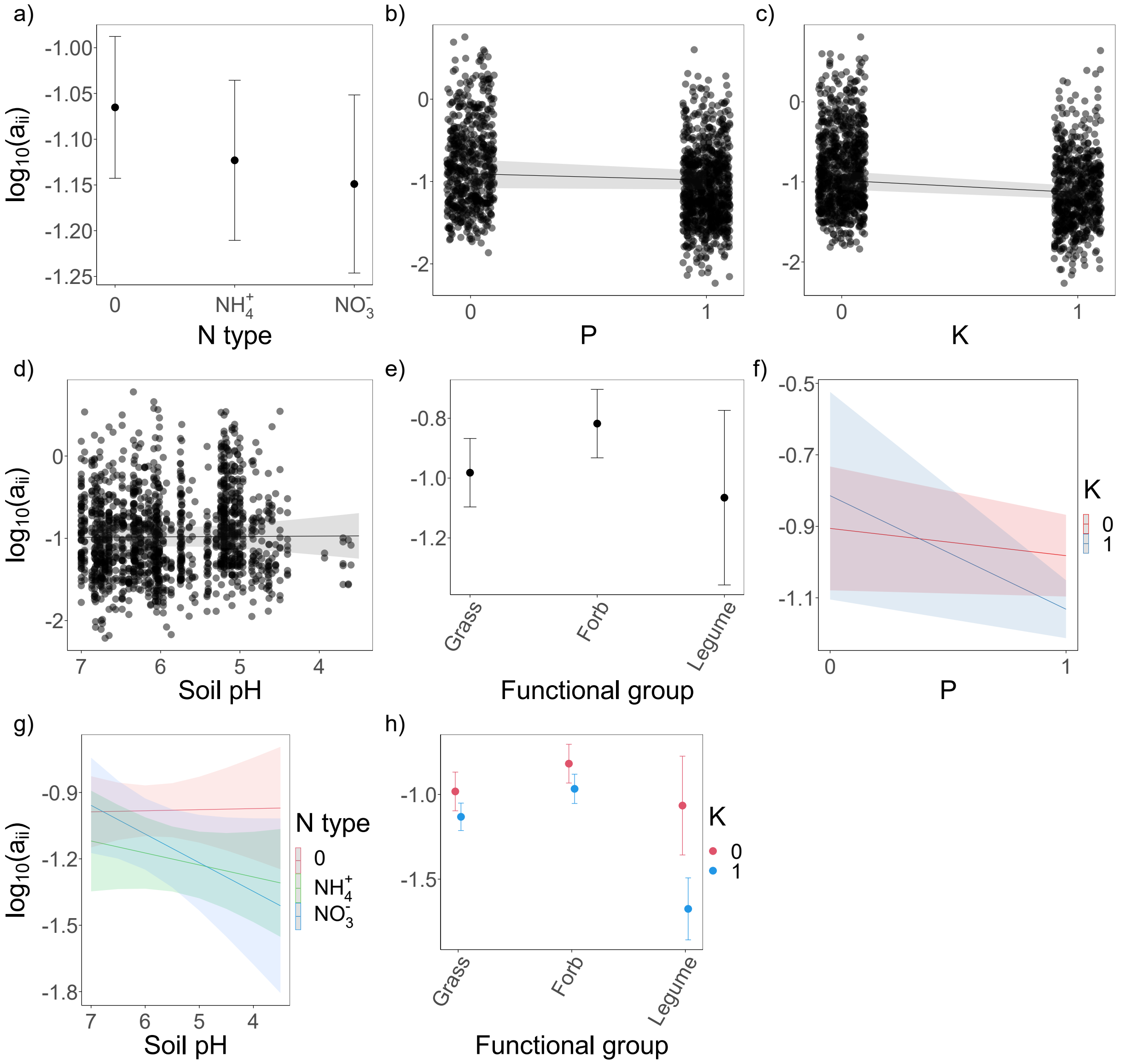

**Fig. S5.5**

Interspecific competition ($a_{ij}$)

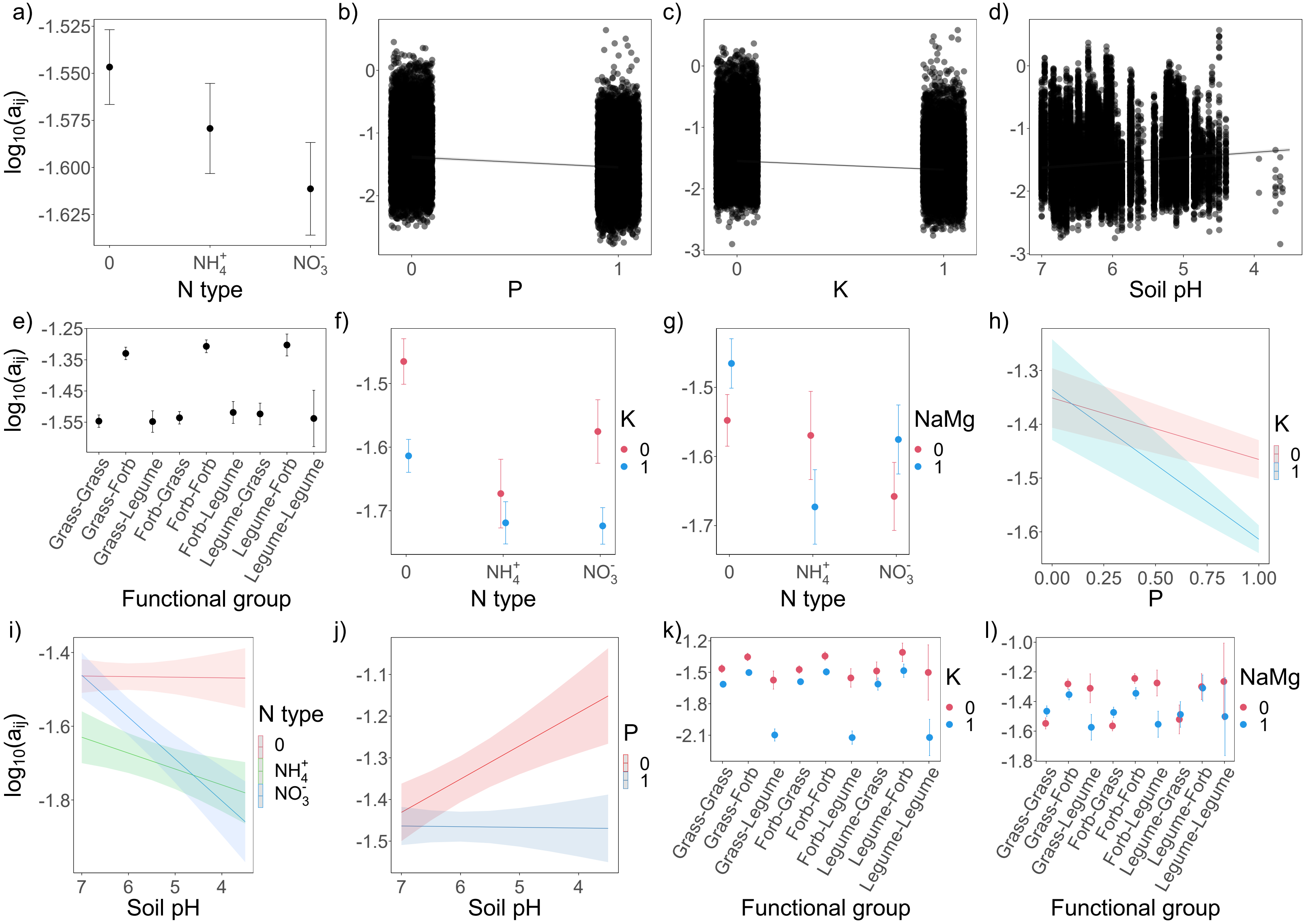

**Table S5** Results of linear models testing the effects of N withholding (N WH: 0, 1), N type ($NH_4^+$, $NO_3^-$), soil pH, functional group (FG) and the interactions between N withholding and functional group and N type on intrinsic population growth rate ($r$), niche difference (ND: $1-\sqrt{\frac{\alpha_{ij}\alpha_{ji}}{\alpha_{ii}\alpha_{jj}}}$), competitive difference (CD: $\sqrt{\frac{\alpha_{jj}\alpha_{ji}}{\alpha_{ii}\alpha_{ij}}}$), and per-capita intraspecific and interspecific competition ($a_{ii}$ and $a_{ij}$). Note that all the variables except for $r$ and ND are $\log_{10}$-transformed to meet the assumptions for residuals. Up and down arrows beside significant (p<0.05, in bold) or marginally significant (0.05⩽p<0.1, in italic) p values indicate positive and negative effects respectively.

| **Variables** | | $\boldsymbol{r}$ | | **ND** | | **CD** | | $\boldsymbol{a_{ii}}$ | | $\boldsymbol{a_{ij}}$ | |
|---|---|---|---|---|---|---|---|---|---|---|---|
| | df | $\chi^2$ | p | $\chi^2$ | p | $\chi^2$ | p | $\chi^2$ | p | $\chi^2$ | p |
| N WH | 1 | 5.6 | **0.018**↑ | 1.23 | 0.267 | 0.03 | 0.858 | 8.57 | **0.003**↑ | 91.9 | **<0.001**↑ |
| N type | 1 | 15.5 | **<0.001** | 0.00 | 0.991 | 0.80 | 0.369 | 0.67 | 0.412 | 8.0 | **0.005** |
| pH | 1 | 19.8 | **<0.001**↑ | 6.87 | **0.009**↓ | 4.87 | **0.027**↓ | 2.40 | 0.122 | 16.1 | **<0.001**↓ |
| FG | 2/5/8 | 13.6 | **0.001** | 4.20 | 0.521 | 8.32 | 0.403 | 18.25 | **<0.001** | 201.1 | **<0.001** |
| N WH: N type | 1 | 3.6 | *0.058* | 3.55 | *0.059* | 0.03 | 0.860 | 0.05 | 0.816 | 5.2 | **0.022** |
| N WH: FG | 2/4/7 | 8.5 | **0.014** | 1.71 | 0.789 | 8.95 | 0.256 | 3.25 | 0.197 | 37.2 | **<0.001** |

**Table S6** Competitive hierarchy of all species *explicitly* fitted in Lotka-Volterra competition models with time-series data in each of the plots with 15 different treatments of nutrient addition (see Box S1 in Appendix S1 for the details of treatments, and also Appendix S2 for the details of model fitting). In the subplot (SP), the "a", "b", "c" and "d" indicate quarter-plots with four different levels of soil pH treatments (soil pH: 7, 6, 5 and unlimed, respectively) in the "recent" subset (1965-2000), and the "U" and "L" indicate the unlimed and limed half-plots respectively in the "longul" subset (1856-2000). Note that all of the U subplots with N added as $(NH_4)_2SO_4$ in the "longul" subset are dropped because of the poor model fitting and cross-validation. The SR indicates the number of *explicitly* fitted species and note that the number of ranked species might be lower than SR because a few species are additionally dropped because of the poor model fitting and cross-validation. Competitive hierarchy is constructed from competitive difference ($\sqrt{\frac{\alpha_{jj}\alpha_{ji}}{\alpha_{ii}\alpha_{ij}}}$) that measures the overall asymmetry within both intraspecific and interspecific competition. Species $i$ is overall more competitive than species $j$ if $\sqrt{\frac{\alpha_{jj}\alpha_{ji}}{\alpha_{ii}\alpha_{ij}}} > 1$, the intraspecific and interspecific competitive effects on species $j$ ($\alpha_{jj}\alpha_{ji}$) are overall greater than those on species $i$ ($\alpha_{ii}\alpha_{ij}$). For abbreviation of species names, see Table S1 in Appendix S1.

| Treatment (plot) | SP | SR | Competitive hierarchy |
|---|---|---|---|
| CK (i.e. Nil) (2/2) | a | 24 | *Leo.his > Tri.rep > Ant.odo > Dac.glo > Tri.pra > Poa.pra > Cen.nig > Luz.cam > Pla.lan > Hel.pub > Lot.cor > Ran.acr > Agr.cap > Hol.lan > Bri.med > Ran.bul > San.min > Tra.pra > Aju.rep > Fes.rub > Con.maj > Lat.pra > Pim.sax > Ach.mil* |
| | b | 20 | *Poa.pra > Cer.fon > Pla.lan > Agr.cap > Hol.lan > Tri.pra > Hel.pub > Bri.med > Cen.nig > Dac.glo > Leo.his > Luz.cam > Ran.acr > Con.maj > Ach.mil > Ant.odo > Fes.rub > San.min > Tar.off > Pim.sax* |
| | c | 18 | *Agr.cap > Ant.odo > Hel.pub > Pla.lan > Dac.glo > Fes.rub > Luz.cam > Leo.his > Ran.acr > Lot.cor > Con.maj > Ach.mil > Car.car > Pim.sax > Cen.nig > Hol.lan > Bri.med > Ran.bul* |
| | d | 14 | *Agr.cap > Luz.cam > Ant.odo > Bri.med > Fes.rub > Ach.mil > Dac.glo > Pla.lan > Car.car > Ran.bul > Cen.nig > Ran.acr > Con.maj > Hel.pub* |
| | U | 23 | *Hol.lan > Tri.fla > Con.maj > Agr.cap > Alo.pra > Bri.med > Dac.glo > Luz.cam > Poa.pra > Ant.odo > Pla.lan > Ach.mil > Fes.rub > Arr.ela > Gal.ver > Car.car > Cen.nig > Hel.pub > Ran.acr > Lot.cor > Aju.rep > Leo.his > Pim.sax* |

| | | | |
|---|---|---|---|
| | L | 25 | *Ant.odo > Tri.rep > Leo.his > Hol.lan > Lat.pra > Hel.pub > Pla.lan > Poa.pra > Tri.pra > Cen.nig > Dac.glo > Luz.cam > Fes.rub > Lot.cor > Agr.cap > Tri.fla > Bri.med > Cer.fon > Con.maj > Ran.acr > Tra.pra > Ach.mil > Alo.pra > Pim.sax > Tar.off* |
| CK (i.e. Nil) (3) | a | 22 | *Pla.lan > Poa.pra > Agr.cap > Bri.med > Hel.pub > Hol.lan > San.min > Cen.nig > Con.maj > Leo.his > Tri.rep > Pim.sax > Ran.bul > Dac.glo > Tri.pra > Luz.cam > Ach.mil > Fes.rub > Ran.acr > Ant.odo > Car.car > Lat.pra* |
| | b | 20 | *Dac.glo > Ant.odo > Luz.cam > Pla.lan > Rum.ace > Leo.his > Poa.pra > Tar.off > Tri.pra > Bri.med > Hel.pub > Ran.acr > Fes.rub > Cen.nig > Cer.fon > Hol.lan > Agr.cap > Ach.mil > Con.maj > San.min* |
| | c | 17 | *Leo.his > Agr.cap > Luz.cam > Ach.mil > Car.car > Pla.lan > Ant.odo > Pil.off > Fes.rub > Bri.med > Cen.nig > Pim.sax > San.min > Con.maj > Lot.cor > Ran.acr > Ran.bul* |
| | d | 13 | *Luz.cam > Ach.mil > Agr.cap > Ant.odo > Fes.rub > Con.maj > Hol.lan > Pla.lan > Bri.med > Dac.glo > Cen.nig > Ran.bul > Leo.his* |
| | U | 23 | *Luz.cam > Tri.pra > Tri.fla > Con.maj > Ant.odo > Lot.cor > Agr.cap > Pla.lan > Ach.mil > Dac.glo > Fes.rub > Rum.ace > San.min > Hel.pub > Poa.pra > Bri.med > Lat.pra > Cen.nig > Hol.lan > Leo.his > Pim.sax > Car.car > Tar.off* |
| | L | 28 | *Ant.odo > Lat.pra > Luz.cam > Tri.pra > Pim.sax > Pla.lan > Poa.pra > Alo.pra > Dac.glo > Tri.rep > Bri.med > Cer.fon > Con.maj > Agr.cap > Fes.rub > Leo.his > Lot.cor > Rum.ace > Tar.off > Hel.pub > Hol.lan > Ach.mil > Arr.ela > Cen.nig > Tri.fla > Ver.cha > Car.car > San.min* |
| CK (i.e. Nil) (12) | a | 23 | *Lat.pra > Hel.pub > Aju.rep > Leo.his > Ach.mil > Luz.cam > Ant.odo > Bri.med > Tri.pra > Poa.pra > Ran.acr > Cen.nig > Hol.lan > Dac.glo > Fes.rub > Pla.lan > Rum.ace > Tar.off > Tri.rep > Cer.fon > Pim.sax > Agr.cap > Con.maj* |
| | b | 24 | *Bri.med > Ran.acr > Hel.pub > Agr.cap > Hol.lan > Pla.lan > Tri.pra > Ant.odo > Fes.pra > Lot.cor > Leo.his > Car.fla > Ach.mil > Car.car > Cen.nig > Dac.glo > Fes.rub > Tar.off > Cer.fon > Lat.pra > Pim.sax > Tri.rep > Con.maj > Luz.cam* |
| | c | 15 | *Ant.odo > Agr.cap > Bri.med > Luz.cam > Pla.lan > Fes.rub > Leo.his > Ach.mil > Dac.glo > Car.car > Pil.off > Cen.nig > Hol.lan > Con.maj > Ran.acr* |
| | d | 14 | *Ant.odo > Leo.his > Fes.rub > Pla.lan > Agr.cap > Luz.cam > Dac.glo > Pim.sax > Ach.mil > Cen.nig > Car.car > Bri.med > Con.maj > Hol.lan* |

| | | | |
|---|---|---|---|
| | U | 20 | *Ant.odo > Tri.pra > Pla.lan > Lot.cor > Fes.rub > Hel.pub > Hol.lan > Leo.his > Pil.off > Agr.cap > Luz.cam > Pim.sax > Ach.mil > Bri.med > Dac.glo > Car.car > Cen.nig > Con.maj > Ran.acr > Tar.off* |
| N1 (1) | a | 19 | *Fes.rub > Hel.pub > Ach.mil > Luz.cam > Pim.sax > Pla.lan > Dac.glo > Ant.odo > Arr.ela > Hol.lan > Leo.his > Poa.pra > Agr.cap > Cen.nig > Rum.ace > Alo.pra > Con.maj > Tar.off > Ran.acr* |
| | b | 14 | *Ach.mil > Agr.cap > Dac.glo > Pla.lan > Cen.nig > Hol.lan > Pim.sax > Luz.cam > Hel.pub > Tar.off > Fes.rub > Ant.odo > Leo.his > Con.maj* |
| | c | 6 | *Agr.cap > Ant.odo > Fes.rub > Rum.ace > Cen.nig > Luz.cam* |
| | d | 2 | *Agr.cap > Ant.odo* |
| | L | 22 | *Poa.pra > Tri.fla > Agr.cap > Cen.nig > Dac.glo > Hol.lan > Pla.lan > Ach.mil > Ant.odo > Lat.pra > Hel.pub > Pim.sax > Rum.ace > Tar.off > Alo.pra > Lot.cor > Luz.cam > Arr.ela > Fes.rub > Leo.his > Con.maj > Ran.acr* |
| N*1 (17) | a | 17 | *Pla.lan > Leo.his > Bri.med > Fes.rub > Agr.cap > Hel.pub > Lol.per > Cen.nig > Ran.acr > Ach.mil > Alo.pra > Dac.glo > Her.sph > Tar.off > Ant.odo > Hol.lan > Con.maj* |
| | b | 17 | *Leo.his > Pla.lan > Fes.rub > Agr.cap > Ant.odo > Car.car > Hol.lan > Ach.mil > Hel.pub > Dac.glo > Bri.med > Ran.acr > Cen.nig > Con.maj > Alo.pra > Lol.per > Tar.off* |
| | c | 16 | *Agr.cap > Leo.his > Ant.odo > Pla.lan > Ran.acr > Con.maj > Her.sph > Cen.nig > Hol.lan > Fes.rub > Ach.mil > Dac.glo > Alo.pra > Car.car > Hel.pub > Tar.off* |
| | d | 17 | *Cen.nig > Fes.rub > Leo.his > Ant.odo > Pla.lan > Ran.acr > Agr.cap > Car.car > Lol.per > Dac.glo > Hol.lan > Tar.off > Hel.pub > Ach.mil > Alo.pra > Con.maj > Luz.cam* |
| | U | 20 | *Pla.lan > Car.car > Fes.rub > Hol.lan > Cen.nig > Dac.glo > Leo.his > Lol.per > Ant.odo > Agr.cap > Alo.pra > Con.maj > Poa.pra > Cer.fon > Hel.pub > Bri.med > Luz.cam > Rum.ace > Ach.mil > Tar.off* |
| | L | 18 | *Fes.rub > Pla.lan > Agr.cap > Alo.pra > Cer.fon > Leo.his > Rum.ace > Ach.mil > Hol.lan > Hel.pub > Ant.odo > Dac.glo > Tar.off > Bri.med > Cen.nig > Lol.per > Con.maj > Luz.cam* |
| P (4/1) | a | 19 | *Ant.odo > Arr.ela > Leo.his > Agr.cap > Pla.lan > Tri.pra > Ach.mil > Pim.sax > Hel.pub > Poa.pra > Rum.ace > Ran.acr > Hol.lan > Luz.cam > Dac.glo > Cer.fon > Fes.rub > Tri.fla > Cen.nig* |
| | b | 22 | *Ran.acr > Leo.his > Ant.odo > Luz.cam > Fes.rub > Pla.lan > Rum.ace > Tri.pra > Cer.fon > Hol.lan > Tar.off > Ach.mil > Lot.cor > Con.maj > Hel.pub > Arr.ela > Dac.glo > Agr.cap > Pim.sax > Cen.nig > Poa.pra > Tri.fla* |
| | c | 14 | *Hol.lan > Pla.lan > Ant.odo > Agr.cap > Luz.cam > Fes.rub > Leo.his > Cen.nig > Con.maj > Hel.pub > Rum.ace > Ach.mil > Ran.acr > Dac.glo* |

| | | | |
|---|---|---|---|
| | d | 13 | *Ran.acr > Pla.lan > Agr.cap > Luz.cam > Ant.odo > Dac.glo > Hol.lan > Leo.his > Ach.mil > Con.maj > Rum.ace > Fes.rub > Cen.nig* |
| | U | 17 | *Ran.acr > Rum.ace > Luz.cam > Agr.cap > Hel.pub > Hol.lan > Pla.lan > Tri.fla > Dac.glo > Leo.his > Ach.mil > Ant.odo > Con.maj > Fes.rub > Cen.nig > Tri.pra > Lot.cor* |
| | L | 27 | *Ran.acr > Luz.cam > Ant.odo > Lat.pra > Leo.his > Fes.rub > Pla.lan > Ach.mil > Rum.ace > Cer.fon > Hol.lan > Con.maj > Tri.fla > Tri.pra > Lot.cor > Tar.off > Hel.pub > Arr.ela > Pim.sax > Dac.glo > Agr.cap > San.min > Kna.arv > Cen.nig > Poa.pra > Bri.med > Ver.cha* |
| N2P (4/2) | a | 10 | *Fes.rub > Agr.cap > Poa.pra > Rum.ace > Ran.acr > Alo.pra > Ant.odo > Hel.pub > Pla.lan > Tar.off* |
| | b | 6 | *Fes.rub > Agr.cap > Alo.pra > Rum.ace > Ant.odo > Poa.pra* |
| | c | 6 | *Fes.rub > Agr.cap > Poa.pra > Hol.lan > Rum.ace > Ant.odo* |
| | d | 2 | *Ant.odo > Agr.cap* |
| | L | 8 | *Fes.rub > Alo.pra > Agr.cap > Poa.pra > Rum.ace > Ant.odo > Hol.lan > Dac.glo* |
| PNaMg (8) | a | 20 | *Ant.odo > Luz.cam > Rum.ace > Hol.lan > Pla.lan > Poa.pra > Tri.pra > Agr.cap > Bri.med > Dac.glo > Cen.nig > Fes.rub > Ach.mil > Tri.fla > Fes.pra > Hel.pub > Arr.ela > Cer.fon > Leo.his > Ran.acr* |
| | b | 21 | *Leo.his > Poa.pra > Tri.pra > Bri.med > Tri.fla > Cer.fon > Dac.glo > Lot.cor > Ant.odo > Pla.lan > Rum.ace > Agr.cap > Cen.nig > Hel.pub > Ran.acr > Con.maj > Fes.rub > Luz.cam > Ach.mil > Arr.ela > Hol.lan* |
| | c | 13 | *Ant.odo > Agr.cap > Hol.lan > Pla.lan > Ran.acr > Leo.his > Luz.cam > Ach.mil > Fes.rub > Rum.ace > Dac.glo > Con.maj > Cen.nig* |
| | d | 13 | *Pla.lan > Agr.cap > Ant.odo > Leo.his > Cen.nig > Dac.glo > Hol.lan > Con.maj > Fes.rub > Ran.acr > Ach.mil > Luz.cam > Lot.cor* |
| | U | 23 | *Arr.ela > Lat.pra > Tri.fla > Hel.pub > Pla.lan > Rum.ace > Agr.cap > Poa.pra > Ant.odo > Leo.his > Hol.lan > Dac.glo > Cen.nig > Cer.fon > Con.maj > Ran.acr > Fes.rub > Luz.cam > Alo.pra > Lot.cor > Tri.pra > Ach.mil > Tar.off* |
| | L | 27 | *Bri.med > Dac.glo > Lol.per > Poa.pra > Con.maj > Pim.sax > Leo.his > Poa.tri > Tri.pra > Tri.fla > Hel.pub > Kna.arv > Lot.cor > Rum.ace > Ant.odo > Pla.lan > Cer.fon > Cen.nig > Agr.cap > Hol.lan > Ran.acr > Ver.cha > Ach.mil > Alo.pra > Arr.ela > Luz.cam > Fes.rub* |
| PKNaMg | a | 17 | *Lat.pra > Tri.pra > Alo.pra > Tar.off > Rum.ace > Arr.ela > Her.sph > Hol.lan > Pla.lan > Ran.acr > Poa.tri >* |

| | | | |
|---|---|---|---|
| (7) | | | *Ant.syl > Poa.pra > Dac.glo > Tra.pra > Fes.rub > Agr.cap* |
| | b | 18 | *Hol.lan > Tri.pra > Dac.glo > Pla.lan > Ran.acr > Alo.pra > Poa.tri > Tar.off > Agr.cap > Her.sph > Rum.ace > Arr.ela > Ant.odo > Ant.syl > Lat.pra > Cer.fon > Poa.pra > Con.maj* |
| | c | 17 | *Agr.cap > Tar.off > Rum.ace > Alo.pra > Cen.nig > Lat.pra > Fes.rub > Ran.acr > Ant.odo > Dac.glo > Con.maj > Tri.pra > Hol.lan > Ach.mil > Pla.lan > Arr.ela > Poa.pra* |
| | d | 14 | *Rum.ace > Ant.odo > Agr.cap > Alo.pra > Fes.rub > Cen.nig > Hol.lan > Lat.pra > Con.maj > Dac.glo > Ach.mil > Pla.lan > Poa.pra > Luz.cam* |
| | U | 19 | *Tri.fla > Rum.ace > Dac.glo > Lat.pra > Agr.cap > Hel.pub > Hol.lan > Ant.odo > Cen.nig > Lot.cor > Fes.rub > Tri.pra > Con.maj > Alo.pra > Pla.lan > Poa.pra > Arr.ela > Luz.cam > Ach.mil* |
| | L | 22 | *Dac.glo > Hel.pub > Hol.lan > Alo.pra > Pla.lan > Tri.pra > Her.sph > Poa.tri > Ach.mil > Agr.cap > Fes.rub > Tar.off > Arr.ela > Poa.pra > Rum.ace > Tri.fla > Bro.hor > Lat.pra > Ant.odo > Cer.fon > Con.maj > Cen.nig* |
| PKNaMg (15) | a | 20 | *Lat.pra > Tra.pra > Ant.syl > Poa.pra > Tri.pra > Con.maj > Her.sph > Pla.lan > Arr.ela > Tar.off > Hel.pub > Poa.tri > Ach.mil > Ant.odo > Fes.rub > Ran.acr > Agr.cap > Alo.pra > Dac.glo > Rum.ace* |
| | b | 20 | *Arr.ela > Hol.lan > Fes.rub > Con.maj > Rum.ace > Tri.pra > Ach.mil > Ran.acr > Agr.cap > Ant.odo > Lat.pra > Tar.off > Dac.glo > Pla.lan > Her.sph > Poa.pra > Tra.pra > Alo.pra > Ant.syl > Hel.pub* |
| | c | 12 | *Hol.lan > Dac.glo > Agr.cap > Fes.rub > Cen.nig > Ant.odo > Tri.pra > Con.maj > Ach.mil > Pla.lan > Alo.pra > Poa.pra* |
| | d | 10 | *Agr.cap > Pla.lan > Ach.mil > Ant.odo > Cen.nig > Fes.rub > Tri.pra > Tar.off > Dac.glo > Con.maj* |
| | U | 20 | *Luz.cam > Tri.fla > Agr.cap > Tri.pra > Ach.mil > Cen.nig > Fes.rub > Pla.lan > Ant.odo > Rum.ace > Alo.pra > Arr.ela > Dac.glo > Hol.lan > Lat.pra > Poa.pra > Tar.off > Hel.pub > Con.maj > Tra.pra* |
| | L | 19 | *Arr.ela > Fes.rub > Hol.lan > Ach.mil > Lat.pra > Poa.pra > Tri.pra > Rum.ace > Tar.off > Ant.odo > Pla.lan > Poa.tri > Alo.pra > Con.maj > Dac.glo > Hel.pub > Agr.cap > Her.sph > Tra.pra* |
| N2PNaMg (10) | a | 11 | *Agr.cap > Fes.rub > Ran.acr > Poa.pra > Rum.ace > Ant.odo > Arr.ela > Hol.lan > Pla.lan > Alo.pra > Tar.off* |
| | b | 8 | *Alo.pra > Fes.rub > Agr.cap > Rum.ace > Ant.odo > Poa.pra > Arr.ela > Hol.lan* |
| | c | 6 | *Fes.rub > Rum.ace > Agr.cap > Ant.odo > Hol.lan > Poa.pra* |
| | d | 3 | *Ant.odo > Hol.lan > Agr.cap* |

| | | | |
|---|---|---|---|
| | L | 10 | *Alo.pra > Fes.rub > Arr.ela > Hel.pub > Agr.cap > Dac.glo > Rum.ace > Ant.odo > Poa.pra > Hol.lan* |
| N2KNaMg (18) | a | 12 | *Ant.odo > Leo.aut > Tar.off > Pla.lan > Dac.glo > Fes.rub > Lol.per > Her.sph > Cen.nig > Leo.his > Tra.pra > Agr.cap* |
| | b | 9 | *Agr.cap > Ant.odo > Dac.glo > Fes.rub > Tra.pra > Cen.nig > Pla.lan > Leo.his > Tar.off* |
| | c | 7 | *Agr.cap > Ant.odo > Lol.per > Dac.glo > Fes.rub > Hol.lan > Poa.tri* |
| | d | 2 | *Agr.cap > Ant.odo* |
| | L | 17 | *Poa.pra > Rum.ace > Hol.lan > Ant.odo > Arr.ela > Cen.nig > Fes.rub > Agr.cap > Dac.glo > Hel.pub > Pla.lan > Tra.pra > Ach.mil > Alo.pra > Her.sph > Tar.off > Con.maj* |
| N1PKNaMg (6) | a | 19 | *Tri.pra > Hol.lan > Pla.lan > Arr.ela > Alo.pra > Poa.tri > Tar.off > Tra.pra > Her.sph > Rum.ace > Dac.glo > Ach.mil > Hel.pub > Lat.pra > Poa.pra > Ran.acr > Ant.odo > Con.maj > Fes.rub* |
| | b | 14 | *Tar.off > Alo.pra > Rum.ace > Ant.odo > Hol.lan > Lat.pra > Ran.acr > Tri.pra > Pla.lan > Dac.glo > Con.maj > Poa.pra > Her.sph > Poa.tri* |
| N*1PKNaMg (16) | a | 17 | *Rum.ace > Ant.syl > Her.sph > Arr.ela > Dac.glo > Pla.lan > Poa.tri > Ran.acr > Tri.pra > Ant.odo > Tar.off > Ach.mil > Fes.rub > Lat.pra > Agr.cap > Con.maj > Poa.pra* |
| | b | 15 | *Alo.pra > Ant.odo > Fes.rub > Tri.pra > Ant.syl > Her.sph > Dac.glo > Ran.acr > Pla.lan > Agr.cap > Con.maj > Poa.tri > Arr.ela > Tar.off > Rum.ace* |
| | c | 16 | *Alo.pra > Pla.lan > Con.maj > Agr.cap > Her.sph > Ant.odo > Fes.rub > Arr.ela > Ran.acr > Lat.pra > Dac.glo > Tri.pra > Ach.mil > Hol.lan > Tar.off > Hel.pub* |
| | d | 14 | *Ach.mil > Lat.pra > Agr.cap > Ant.odo > Lol.per > Fes.rub > Ran.acr > Con.maj > Hol.lan > Tar.off > Tri.pra > Her.sph > Pla.lan > Dac.glo* |
| | U | 18 | *Ach.mil > Lat.pra > Poa.tri > Agr.cap > Ant.odo > Alo.pra > Lol.per > Hol.lan > Ran.acr > Con.maj > Fes.rub > Hel.pub > Tar.off > Tri.pra > Dac.glo > Poa.pra > Rum.ace > Pla.lan* |
| | L | 21 | *Rum.ace > Ant.syl > Her.sph > Arr.ela > Alo.pra > Dac.glo > Poa.tri > Ran.acr > Pla.lan > Tra.pra > Tri.pra > Con.maj > Ant.odo > Tar.off > Ach.mil > Bro.hor > Agr.cap > Lat.pra > Fes.rub > Poa.pra > Hel.pub* |
| N2PKNaMg (9/2) | a | 13 | *Arr.ela > Her.sph > Hol.lan > Poa.tri > Tar.off > Ant.syl > Lat.pra > Rum.ace > Alo.pra > Dac.glo > Fes.rub > Poa.pra > Ant.odo* |
| | b | 11 | *Ant.syl > Alo.pra > Arr.ela > Dac.glo > Hol.lan > Poa.pra > Con.maj > Her.sph > Rum.ace > Ant.odo > Tar.off* |

| | | | |
|---|---|---|---|
| | c | 9 | *Agr.cap > Hol.lan > Fes.rub > Ant.odo > Rum.ace > Tri.pra > Arr.ela > Her.sph > Poa.pra* |
| | d | 3 | *Agr.cap > Ant.odo > Hol.lan* |
| | L | 12 | *Alo.pra > Arr.ela > Agr.cap > Ant.odo > Hol.lan > Poa.pra > Fes.rub > Tar.off > Lat.pra > Rum.ace > Dac.glo > Her.sph* |
| (N2)PKNaMg (9/1) | a | 15 | *Arr.ela > Ant.odo > Hol.lan > Dac.glo > Tar.off > Con.maj > Poa.pra > Agr.cap > Tri.pra > Fes.rub > Her.sph > Rum.ace > Ant.syl > Alo.pra > Lat.pra* |
| | b | 13 | *Alo.pra > Her.sph > Tar.off > Agr.cap > Dac.glo > Hol.lan > Ant.odo > Arr.ela > Con.maj > Fes.rub > Poa.pra > Rum.ace > Lat.pra* |
| | c | 7 | *Agr.cap > Ant.odo > Fes.rub > Her.sph > Poa.pra > Hol.lan > Rum.ace* |
| | d | 3 | *Ant.odo > Agr.cap > Hol.lan* |
| N*2PKNaMg (14/2) | a | 10 | *Poa.tri > Arr.ela > Ant.syl > Rum.ace > Tar.off > Dac.glo > Alo.pra > Fes.rub > Ran.acr > Poa.pra* |
| | b | 11 | *Poa.tri > Alo.pra > Ant.syl > Arr.ela > Poa.pra > Ran.acr > Tar.off > Fes.rub > Her.sph > Rum.ace > Dac.glo* |
| | c | 12 | *Poa.tri > Alo.pra > Arr.ela > Rum.ace > Poa.pra > Dac.glo > Her.sph > Bro.hor > Fes.rub > Ran.acr > Tar.off > Ant.syl* |
| | d | 13 | *Arr.ela > Alo.pra > Her.sph > Poa.tri > Hol.lan > Ran.acr > Tar.off > Ant.syl > Fes.rub > Poa.pra > Rum.ace > Bro.hor > Dac.glo* |
| | U | 12 | *Arr.ela > Poa.tri > Alo.pra > Bro.hor > Hol.lan > Rum.ace > Agr.cap > Poa.pra > Tar.off > Dac.glo > Lat.pra > Fes.rub* |
| | L | 16 | *Lat.pra > Tri.fla > Alo.pra > Poa.tri > Arr.ela > Ant.syl > Bro.hor > Dac.glo > Poa.pra > Ran.acr > Fes.rub > Rum.ace > Agr.cap > Her.sph > Tar.off > Ant.odo* |
| (N*2)PKNaMg (14/1) | a | 17 | *Bro.hor > Pla.lan > Tar.off > Arr.ela > Rum.ace > Poa.tri > Her.sph > Lat.pra > Ran.acr > Ant.syl > Fes.rub > Hel.pub > Alo.pra > Poa.pra > Tri.pra > Dac.glo > Con.maj* |
| | b | 17 | *Ant.syl > Rum.ace > Tra.pra > Arr.ela > Bro.hor > Tri.pra > Alo.pra > Fes.rub > Poa.pra > Her.sph > Ant.odo > Dac.glo > Hol.lan > Tar.off > Ran.acr > Pla.lan > Poa.tri* |
| | c | 17 | *Bro.hor > Lat.pra > Poa.pra > Ran.acr > Pla.lan > Tar.off > Ant.syl > Agr.cap > Dac.glo > Hol.lan > Rum.ace > Ant.odo > Poa.tri > Fes.rub > Her.sph > Arr.ela > Alo.pra* |

| | | | |
|---|---|---|---|
| | d | 17 | *Ant.odo > Tar.off > Agr.cap > Fes.rub > Pla.lan > Dac.glo > Hol.lan > Ran.acr > Rum.ace > Poa.tri > Lat.pra > Tri.pra > Her.sph > Alo.pra > Poa.pra > Arr.ela > Bro.hor* |
| N3PKNaMg (11/1) | a | 10 | *Alo.pra > Arr.ela > Poa.tri > Rum.ace > Dac.glo > Hol.lan > Her.sph > Tar.off > Ant.syl > Poa.pra* |
| | b | 9 | *Alo.pra > Arr.ela > Hol.lan > Dac.glo > Her.sph > Rum.ace > Tar.off > Ant.syl > Poa.pra* |
| | c | 6 | *Alo.pra > Poa.pra > Ant.odo > Dac.glo > Agr.cap > Hol.lan* |
| | d | 1 | *Hol.lan* |
| | L | 10 | *Alo.pra > Fes.rub > Ant.odo > Her.sph > Hol.lan > Poa.pra > Tar.off > Dac.glo > Agr.cap > Rum.ace* |
| N3PKNaMgSi (11/2) | a | 7 | *Alo.pra > Ant.syl > Arr.ela > Hol.lan > Poa.tri > Dac.glo > Her.sph* |
| | b | 4 | *Alo.pra > Hol.lan > Her.sph > Dac.glo* |
| | c | 4 | *Alo.pra > Hol.lan > Agr.cap > Poa.pra* |
| | d | 1 | *Hol.lan* |
| | L | 9 | *Alo.pra > Rum.ace > Tar.off > Agr.cap > Ant.odo > Poa.pra > Her.sph > Hol.lan > Dac.glo* |

**Table S7** Results of generalized linear models testing the effects of the changes (due to various treatments of nutrient addition and soil pH) in intrinsic population growth rate ($r$), niche difference (ND: $1-\sqrt{\frac{\alpha_{ij}\alpha_{ji}}{\alpha_{ii}\alpha_{jj}}}$), absolute competitive difference (CD: $\sqrt{\frac{\alpha_{jj}\alpha_{ji}}{\alpha_{ii}\alpha_{ij}}}$), and their interactions on species richness. Note that all the response and explanatory variables are plot-level averages, and species richness is averaged from 1991 to 2000. Absolute competitive differences (abs CD) are absolute values of $\log_{10}$-transformed competitive differences, which are supposed to reflect absolute competitive asymmetries driving competitive exclusion. Up and down arrows beside significant (p<0.05, in bold) or marginally significant (0.05⩽p<0.1, in italic) p values indicate positive and negative effects respectively.

| **Variables** | **Species richness** | | |
|---|---|---|---|
| | df | $\chi^2$ | p |
| Intrinsic population growth rate ($r$) | 1 | 46.42 | **<0.001**↑ |
| Niche difference (ND) | 1 | 4.63 | **0.031**↑ |
| Absolute competitive difference (abs CD) | 1 | 53.77 | **<0.001**↓ |
| $r$: ND | 1 | 20.27 | **<0.001** |
| $r$: abs CD | 1 | 15.72 | **<0.001** |
| ND: abs CD | 1 | 11.73 | **<0.001** |

### Appendix S4 Results of structural niche and competitive differences

Structural niche and competitive differences (SND and SCD) have been recently proposed to understand multispecies coexistence (8). In theory, SND reflects stabilization potential as they define the feasibility domain where a multispecies community is feasible and stable. Mathematically, SND is defined as the solid angel ($\Omega$) of the algebraic cone bounded by the column vectors of the competitive coefficient matrix ($\boldsymbol{\alpha}$) of multiple species. SCD corresponds to the angle ($\theta$) between the centroid of the cone describing the feasibility domain ($\boldsymbol{r_c}$) and the observed vector of intrinsic growth rates ($\boldsymbol{r}$). The $\boldsymbol{r_c}$ is calculated by the column vectors of $\boldsymbol{\alpha}$ ($\boldsymbol{V_i}$). To fit into our models, the $\boldsymbol{\alpha}$ in Saavedra's framework is replaced with $\boldsymbol{\beta}$ that is $\boldsymbol{\alpha}$ multiplied by $\boldsymbol{R}$, a diagonal matrix with $\boldsymbol{r}$ as the diagonal. Note, although SND and SCD are multispecies measures, in fact, they still only account for species interactions that are essentially pairwise (8). In other words, Saavedra*, et al.* (8) includes intransitive competition ("paper-rock-scissors" game) but not *higher-order interactions (HOIs)* (9, 10). More details about SND and SFD can be found in Saavedra*, et al.* (8). In our study, SND was strongly negatively correlated with species richness ($\rho$=-0.62, p<0.001). Figs S6 and S7 show the changes of SND and SCD across the 15 different treatments of nutrient addition in which one plot has one value for SND or SCD. Similar to the *pairwise* niche difference (Table S4 in Appendix S3), SND slightly increased with N type (Table S8).

$$\boldsymbol{\beta} = \boldsymbol{R} \cdot \boldsymbol{\alpha}$$

$$\Omega(\boldsymbol{\beta}) = \frac{|\det(\boldsymbol{\beta})|}{\sqrt[n]{\frac{\pi}{2}}} \int \dots \int_{\boldsymbol{R_n} \leq 0} e^{-\boldsymbol{x}^T \boldsymbol{\beta}^T \boldsymbol{\beta} \boldsymbol{x}} \mathrm{d}\boldsymbol{x}$$

$$\boldsymbol{r_c} = \frac{1}{n}\left(\frac{\boldsymbol{V_1}}{\|\boldsymbol{V_1}\|} + \frac{\boldsymbol{V_2}}{\|\boldsymbol{V_2}\|} + \dots + \frac{\boldsymbol{V_n}}{\|\boldsymbol{V_n}\|}\right)$$

$$\theta = arccos\frac{\boldsymbol{r} \cdot \boldsymbol{r_c}}{\|\boldsymbol{r}\| \cdot \|\boldsymbol{r_c}\|}$$

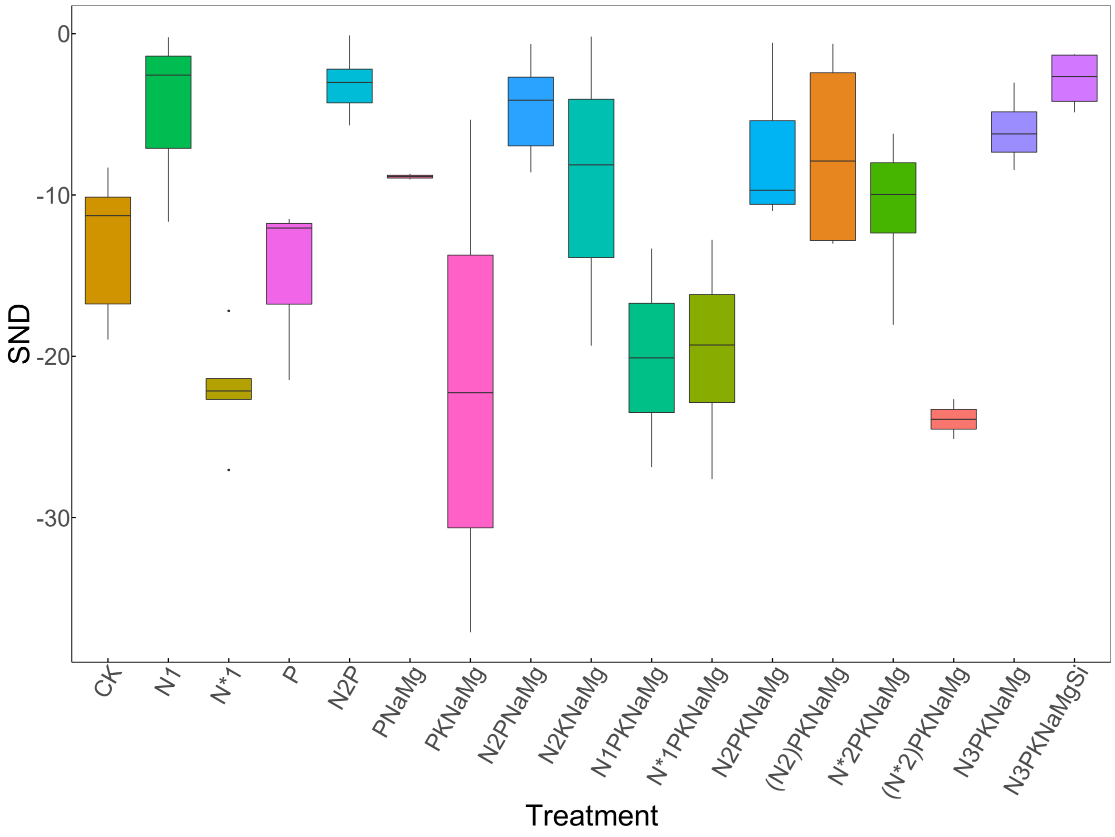


**Fig. S6** Structural niche differences (SND) across the 15 different treatments of nutrient addition (see Box S1 in Appendix S1 for the details of treatments). Please note (N2)PKNaMg and (N*)2PKNaMg indicate N addition is withheld in one half of N2PKNaMg and N*2PKNaMg respectively since 1990.

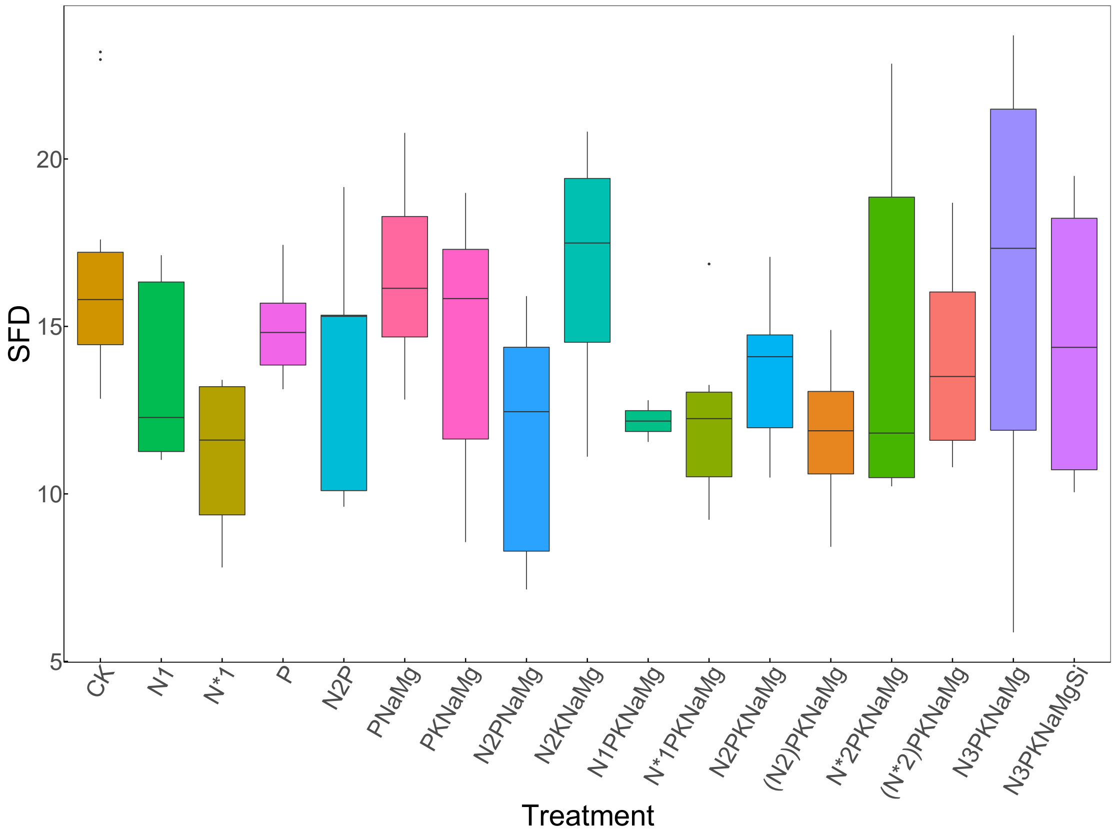


**Fig. S7** Structural competitive differences (SCD) across the 15 different treatments of nutrient addition (see Box S1 in Appendix S1 for the details of treatments). Please note (N2)PKNaMg and (N*)2PKNaMg indicate N addition is withheld in one half of N2PKNaMg and N*2PKNaMg respectively since 1990.

**Table S8** Results of linear models testing the effects of N type (0, $NH_4$, $NO_3$), P (0, 1), K (0, 1), NaMg (0, 1), soil pH and their interactions on structural niche and competitive differences (SND and SCD). For SND, please note that species richness (SR) is also included as a covariate because of its strong negative correlation with SND ($\rho$=-0.62, p<0.001). Up and down arrows beside significant (p<0.05, in bold) or marginally significant (0.05⩽p<0.1, in italic) p values indicate the positive and negative effects respectively.

| **Variables** | | **SND** | | **SCD** | |
|---|---|---|---|---|---|
| | df | $\chi^2$ | p | $\chi^2$ | p |
| N type | 2 | 5.76 | *0.056*↑ | 2.08 | 0.353 |
| P | 1 | 0.56 | 0.454 | 0.15 | 0.701 |
| K | 1 | 1.60 | 0.205 | 0.55 | 0.459 |
| NaMg | 1 | 0.07 | 0.787 | 0.15 | 0.698 |
| pH | 1 | 2.12 | 0.145 | 0.32 | 0.570 |
| SR | 1 | 5.90 | **0.015**↓ | 1.04 | 0.309 |
| N type: P | 1 | 0.30 | 0.587 | 0.32 | 0.571 |
| N type: K | 1 | 4.69 | **0.030** | 4.12 | **0.042** |
| N type: NaMg | 1 | 1.63 | 0.202 | 2.83 | *0.093* |
| P: K | 1 | 0.84 | 0.360 | 0.96 | 0.326 |
| pH: N type | 2 | 3.36 | 0.187 | 1.00 | 0.608 |
| pH: P | 1 | 0.73 | 0.394 | <0.01 | 0.960 |
| pH: K | 1 | 1.12 | 0.291 | 0.07 | 0.788 |
| pH: NaMg | 1 | 0.19 | 0.664 | 3.91 | **0.048** |
